# Insights into the regulatory mechanisms of *Clostridioides difficile* biofilm formation

**DOI:** 10.1101/2021.02.19.431970

**Authors:** Anthony M. Buckley, Duncan Ewin, Ines B. Moura, Mark H. Wilcox, Gillian R. Douce

**Author notes:** Corresponding author: Dr Gillian Douce, Institute of Infection, Immunity and Inflammation, University of Glasgow, Glasgow, U.K. G12 8TA.

## Abstract

Mucosal biofilms play an important role in intestinal health; however, the mucosal bacterial community has been implicated in persistent infections. *Clostridioides difficile* is an important nosocomial pathogen, with an unacceptable high rate of recurrence following antibiotic treatment. As *C. difficile* is a known biofilm producer, a property which may contribute to this suboptimal therapeutic response, we have investigated the transcriptional changes and regulatory pathways during the transition from planktonic to biofilm mode of growth. Widespread metabolic reprogramming during biofilm formation was detected, characterised by an increased usage of glycine metabolic pathways to yield key metabolites, which are used for energy production and synthesis of short chain fatty acids. We detected the expression of 107 small non-coding RNAs that appear to, in some part, regulate these pathways; however, 25 of these small RNAs were specifically expressed during biofilm formation, indicating they may play a role in regulating biofilm-specific genes. Similar to *Bacillus subtilis*, biofilm formation is a multi-regulatory process and SinR negatively regulates biofilm formation independently of other known mechanisms. This comprehensive analysis furthers our understanding of biofilm formation in *C. difficile*, identifies potential targets for anti-virulence factors, and provides evidence of the link between metabolism and virulence traits.

## INTRODUCTION

*Clostridium difficile* is the most common infective cause of antibiotic associated diarrhoea, causing a huge burden on health care facilities around the world. *C. difficile* infection (CDI) can manifest as mild, self-limiting diarrhoea to pseudomembranous colitis, colonic perforation and death. The majority of symptoms are due to the production of two large toxins, TcdA and TcdB^1^, the actions of which cause the loss of epithelial barrier integrity^2^. The proportion of CDI cases that recur after primary antibiotic therapy is 20-30%^3^, with most representing relapses due to the original strain. This suggests that *C. difficile* can occupy a protective niche, which can play a role in long-term persistence/colonisation *in vivo*. Indeed, we have recently shown that sessile *C. difficile* cells contribute towards recurrent disease^4^.

The term ‘biofilm’ describes microbial populations enclosed in a self-produced matrix adherent to each other and/or surfaces. Biofilms are ubiquitous in nature and can have substantial benefits/consequences depending on the environment and composition of the biofilm. Within the human gastrointestinal tract (GI tract), there is growing evidence that the microbiota can exist in two forms: luminal planktonic bacteria and mucosal sessile bacteria^5^, where the formation of biofilms can affect the function of the intestinal microbiome and its interactions with the host^6^. In healthy individuals, the microbiota exists in a mutualistic state with the host, i.e. the host can use bacterial fermentation products of food as an energy source, and bacteria are provided with favourable conditions for growth. Perturbation of this mutualistic relationship, e.g. following administration of antibiotics, causes profound effects on the status quo of the epithelial layer, the metabolic functions of the gut^7^ and eliminates the protective colonisation barrier that allows colonisation or outgrowth of potentially pathogenic bacteria, such as *C. difficile*^8^.

*C. difficile* can form a mono-species biofilm^9^ or associate with an existing poly-species biofilm *in vitro* and *in vivo*^4,10,11^. Strong evidence exists to suggest that sessile *C. difficile* biofilms exist as a heterologous population, made up of vegetative cells, spores and micro-aggregates, all encased in a glycan-rich extracellular matrix, composed mostly of polysaccharide PS-II^12,13^. Poquet *et al*.,^14^ characterised the genetic pathways present in strain 630Δ*erm* during mid-stage biofilm development (72 hr growth); sessile cells were found to have a different metabolic profile, remodelled cell wall and envelope and display different cell surface organelles compared with planktonic cells.

The genetic factors that control the switch of planktonic motile *C. difficile* cells to sessile non-motile cells are largely unknown; however, several cell-surface structures, such as Type IV pili (T4P)^15,16^, flagella^17,18^ and S-layer^12,19^ have been shown to play a role in biofilm formation (reviewed in^20^). Some of these cell surface macromolecules can be genetically controlled by riboswitches in response to the secondary cell messenger Bis-(3’-5’)-cyclic dimeric guanosine monophosphate (c-di-GMP). High c-di-GMP levels increase the expression of the T4P loci^21^, by interacting with a type II riboswitch (Cdi2_4)^22^, whilst repressing the expression of the major flagella operon *flgB*^23^ via a type I riboswitch (Cdi1_3)^22,24^. In this way, a single signalling molecule can help coordinate cell lifestyle differentiation within a population.

In *Bacillus subtilis*, there are several genetic pathways that promote biofilm formation, one through post-transcriptional phosphorylation of Spo0A, which in turn regulates the SinI-SinR locus, allowing activation of matrix production genes (reviewed in^25^). SinR represses the genetic loci needed for biofilm formation, but *sinI* expression overcomes this repression, leading to biofilm formation. *C. difficile* genomes contain a two gene locus similar to the genetic arrangement in *B. subtilis*, corresponding to a homologue of SinR (CD2214 in 630 and CDR20291_2121 in R20291). Another gene at this locus, CDR20291_2122, has been reported; however, there is a discrepancy in the literature whether this second gene is a homologue of SinI or if this is an extra SinR homologue (termed SinR’)^26–28^. *C. difficile* SinR and SinR’ are master regulators that play a role in regulating biofilm formation, motility, sporulation and toxin production^14,27,28^. Inhibition of flagella rotation is another genetic pathway that promotes biofilm formation in *B. subtilis*, through phosphorylation of the pleotropic regulator, DegU^29^. A similar phenotype has been seen in *C. difficile*, where inhibition of flagella rotation, either through deletion of flagella motor (*motB*) or preventing flagella glycosylation, resulted in increased biofilm formation^17,30^, although the underlying mechanisms for this are unclear.

In our study, we define the genetic pathways during early stages of biofilm development in the clinical *C. difficile* ribotype 027 strain R20291. Through defined chromosomal mutations, we explore the role of CDR20291_2121 during biofilm formation, and the interplay of different regulatory pathways involved biofilm formation.

## MATERIALS AND METHODS

### Bacterial strains and growth conditions

Bacterial strains used in this study are listed in **Supplemental Table S1**. *Escherichia coli* TOP10 (Invitrogen, U.K.), used as a cloning host and *E. coli* CA434^31^ as a conjugal donor were grown aerobically on Luria Bertani medium supplemented with ampicillin (100 µg/ml^-1^) or chloramphenicol (15 µg/ml^-1^) when required. *C. difficile* strains were routinely grown on either CCEY agar plates supplemented with cefoxitin-cycloserine and 5% horse blood or Brain-heart infusion (BHI) agar in an anaerobic workstation (Don Whitley Scientific Ltd, U.K.) at 37 °C. Brain-heart infusion (BHI) broth supplemented with yeast extract and cycteine was routinely used to grow *C. difficile* strains. Plasmids used in this study are shown in **Supplemental Table S1 and Figure S1**.

### Biofilm growth assay

Sterilised 13 mm glass cover slips were placed at the bottom of the well of a 24-well tray. Quadruple 2 ml liquid BHISC cultures were pre-reduced and inoculated (1:10) with overnight cultures of *C. difficile* strains and incubated anaerobically for 3 days. The glass cover slips were thrice washed with PBS and 1 % filter sterilised crystal violet (CV) used to visualise the biofilms. Excess CV was removed by thrice washing the glass cover slip with PBS and using 100 % methanol to dissolve the remaining CV. 1:10 serial dilutions of these cultures were made in PBS and the OD_595_ measured in a Tecan spectrophotometer (Infinite Pro 200, Tecan). A media only negative control was included in all experiments and used to adjust the background CV absorbance. All assays were performed with a minimum of three biological repeats and four technical repeats. A student’s T test was performed to determine if there were differences in biofilm formation between mutant strains and wild-type. To enumerate the total viable counts and the spore cells, 3-day-old biofilms were washed with pre-reduced PBS and mechanically disrupted before 1:10 serial dilutions were made and plated onto CCEY. Spores were enumerated by ethanol shock method before 1:10 serial dilutions made and plated onto CCEY as before.

### Scanning electron microscopy

Biofilm cultures were grown as previously described but for 12 hr, except Thermanox® coverslips were used instead of glass coverslips. Biofilms were fixed and prepared as described by Goulding *et al*,^32^.

### Cytotoxin assay

Supernatant from planktonic grown cultures was centrifuged (14,000 rpm, 10 min, 4°C) and filter sterilised by filtration through 0.22 µM syringe filters. Biofilm cultures were grown washed as above, and then 500 µl sterile PBS added and the biofilm mechanically disrupted. 10-fold serial dilutions of planktonic and biofilm supernatants were applied to culture Vero cells and cytotoxicity levels determined as previously described^10^. Cytotoxin titers were correlated to an arbitrary log_10_ scale and expressed as relative units (RUs) at the highest dilution, with approximately 70% cell rounding (i.e., 100, 1RU; 10^−1^, 2RUs; and 10^−2^, 3RUs).

### RNA extraction, DNase treatment and rRNA removal

Biofilms used for RNA-Seq experiments were grown as described above except, 6-well trays with 5 ml BHISC without glass cover slips were used. Cultures were grown for 12 hr anaerobically before the supernatant removed and replaced with pre-reduced RNA*later* (ThermoFisher Scientific) to wash the biofilm and incubated in a further aliquot of RNA*later* for 30 mins and mechanically disrupted. The wells from each tray were pooled and centrifuged at 4,000 rpm for 30 mins at 4 °C and the pellets frozen at -20 °C until extracted. Alongside each biofilm culture, 10 ml from a shaking culture, grown for 12 hr, was treated with RNA*later* and processed in the same way. Three biological replicates were performed for both culture types.

RNA was extracted using the FastRNA® Pro Blue Kit (MP Biosciences) and the FastPrep® 24 system following the manufacturer’s instructions. Genomic DNA was removed from the RNA samples using Turbo DNA-*free*-™ kit (ThermoFisher Scientific) following the manufacturer’s rigorous treatment protocol. RNA integrity and quantity were checked using the Agilent 2100 bioanalyzer, then rRNA depleted using the Ribo-Zero™ rRNA removal for Gram-positive bacteria (Illumina) following the manufacturer’s instructions. Elute RNA was purified by phenol/chloroform extraction and ethanol precipitation and the pellet resuspended in RNase, DNase free water.

### cDNA preparation and sequencing

cDNA library construction for Ion Torrent sequencing was performed using the Ion Total RNA-Seq Kit, following the manufacturer’s instructions and sequences generated via Ion Torrent sequencing. cDNA preparation and sequencing services were provided by the Glasgow Polyomics suite, University of Glasgow.

### Sequencing data analysis

After sequencing, adapters were trimmed by Ion Torrent internal software during the base calling. CLC genomics workbench version 12.0 (Qiagen) was used for quality, alignment with the *C. difficile* R20291 genome (GenBank Accession Number: FN545816.1), normalisation of the reads per kilobase per million mapped reads and for the analysis of differential gene expression, using default settings for each analysis. A gene was considered to be differentially expressed [DE (genes G)] when the false discovery rate (FDR) *p* value was ≤ 0.05 and/or a -2 ≤ fold change ≥ 2 in comparison with shaking cultures was observed.

DEGs were filtered to remove lowly expressed genes (less than 10 reads). Functional and metabolic pathways were inferred using a variety of means; homology to previously characterised *C. difficile* strain 630 proteins, Gene Ontology terms (UniProt for R20291 protein database)^33^, Blastp homology searches (restricted to all *C. difficile* strains) and the Kyoto Encyclopaedia of Genes and Genomes (KEGG) database. sRNAs were identified by aligning the sequencing read map from each replicate with the R20291 gene track and searching for reads mapped outside of the coding regions. Rfam database was used to predict RNA secondary structure and assign putative Rfam family to that sRNA^34^ RNAfold version 2.4.8^35^ was used to predict the secondary structure of the glycine riboswitches, and structures were viewed using Force-directed RNA (forna) webviewer^36^. RNApredator version 2.4.8^37^ was used to predict the mRNA targets of the novel sRNAs.

### RT-PCR confirmation of transcripts

RT-PCR was used to confirm putative sRNA expression detected by RNASeq within biofilms and shaking cultures. RT-PCR was performed using three independent biological repeats of those samples used for Ion Torrent sequencing. Briefly, RNA was extracted, treated with DNase, rRNA depleted and quality controlled as described above. After rRNA depletion, one shaking replicate failed the QC parameters and was discounted from further processing and analysis. cDNA was constructed using SuperScript® First-strand cDNA synthesis system (ThermoFisher Scientific) following manufacturer’s instructions. cDNA across the samples was normalised and specific transcripts amplified using Phusion® high-fidelity PCR master mix (NEB) with primers outlined in **Supplemental Table S2**. Products were visualised by agarose gel electrophoresis.

### C. difficile R20291 mutant construction

Defined chromosomal CDR20291_2121 mutants, either in wild-type or R20291::Tn0241 backgrounds, were constructed using the method described in^38^. Briefly, approximately 1200 bp up- and downstream of CDR_2121 start and stop codons (**Supplemental Figure S1A**), homology arms 1 & 2, respectively, were amplified using primers outlined in **Supplemental Table S2** and Phusion® high-fidelity PCR master mix with GC buffer. The insert was cloned into the allelic exchange vector pMTL82151 using Gibson Assembly® master mix (**Supplemental Table S2**) and transformed into *E. coli* Topo10 cells (**Supplemental Figure S1B**). Once confirmed, the plasmid was transformed into *E. coli* CA434 cells and conjugated with *C. difficile* R20291 as previously described^38^. *C. difficile* Δ*CDR20291_2121* strains were confirmed by PCR and sequencing (**Supplemental figure S1C**). Using this method, 321/339 bp of CDR20291_2121 were deleted leaving the start and stop codons intact, thus preventing polar effects of downstream genes.

### Construction of complementation vectors

The *CDR20291_2121* complementation plasmid was constructed by PCR amplification from 20 bp upstream of the CDR20291_2121 start codon to the stop codon, the fragment cloned into the constitutive expression vector pRPF144^39^, and transformation of *E. coli* Topo10 cells (**Supplemental Figure S2**). Plasmids were sequenced and transformed into *E. coli* CA434 cells for conjugation to *C. difficile* strains.

### Statistical analysis

A student’s T test in Excel (Microsoft office) was used to determine the significant differences in biofilm formation from the R20291 wild-type strain, mutated strains and complemented strains.

## RESULTS

### Characterising the C. difficile strain R20291 biofilm

The ability of *C. difficile* to form a biofilm *in vitro* has been shown previously^12,13^; however, in this study we grew R20291 biofilms, which are more pronounced compared with other strains^12^. Biofilms were grown on glass coverslips to reduce disruption that occurs during the washing steps. After 3 days, *C. difficile* R20291 was able to form a mature biofilm that was encased in a self-produced extracellular matrix (**Figure 1**). This biofilm was composed of vegetative and spore cells (5:1 ratio, respectively); however, scanning electron microscopy (SEM) analysis detected what appeared to be cellular debris within the biofilms as well. It is unclear whether the cellular debris seen was due to natural cell death (possibly associated with nutrient limitation) or due to programmed cell death. A cannibalism phenotype has been described during *B. subtilis* biofilm formation, whereby a proportion of cells secrete a toxin that causes cell death unless that cell is expressing the anti-toxin^40^. *C. difficile* harbours several of these Toxin-Antitoxin (TA) systems that have a putative role in biofilm formation^41,42^. The observation of spores within the biofilm, as seen previously^43^, alongside vegetative cells suggests a heterogenous population whereby cells undergo cell differentiation to perform different functions within a biofilm.

**Figure 1.**
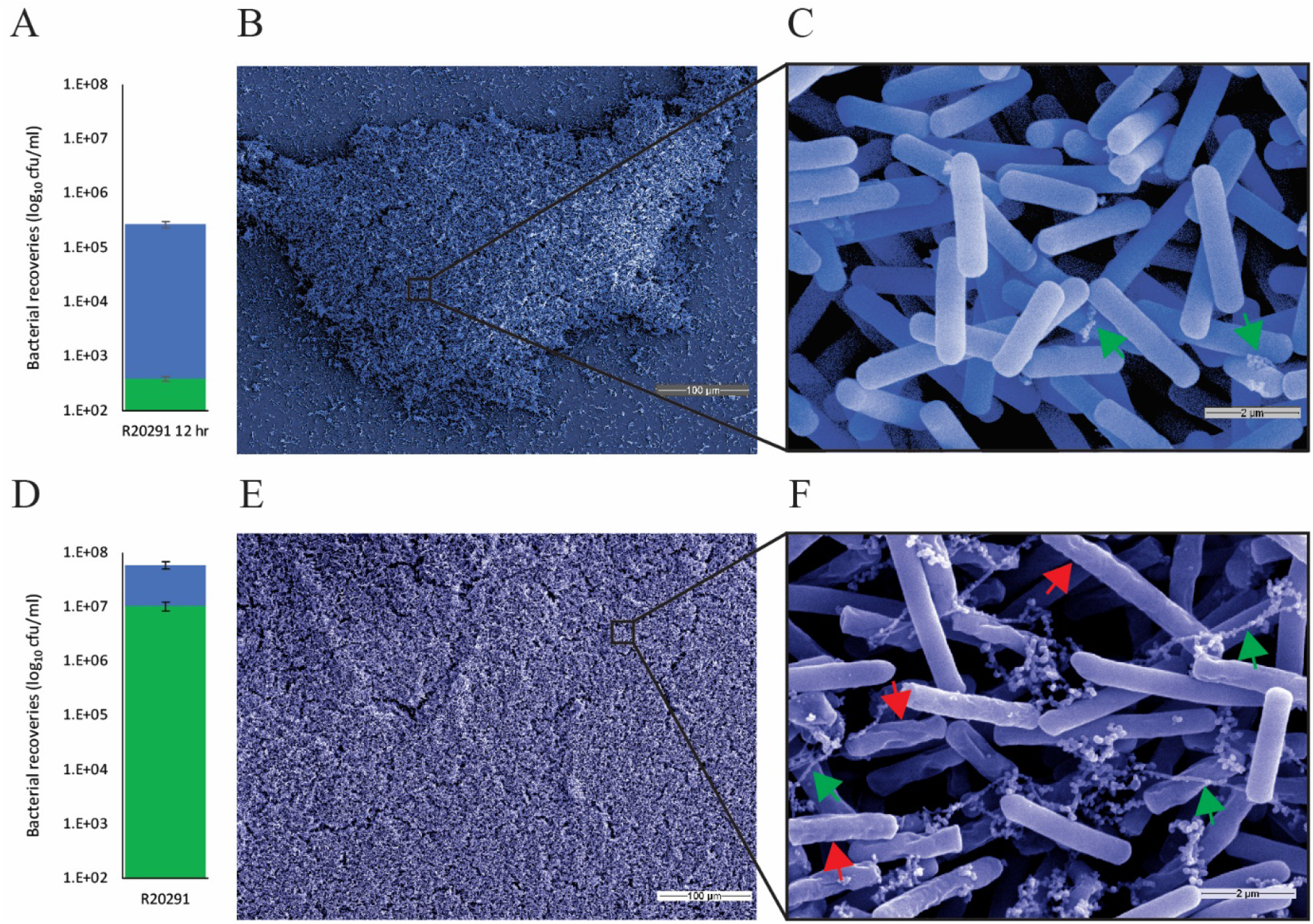
Characterisation of the immature (12 hr – A/B/C) and mature (3 day – E/F/G) *C. difficile* biofilm. (A/E) *C. difficile* biofilm is composed of viable vegetative (blue bar) and spores cells (green bar) (mean ±SEM), forms 3D structures [false-coloured scanning electron micrograph at x200 (B/F) and x10,000 (C/G)], (C/G) where cells are enclosed in a self-produced matrix (greens arrows) and cellular debris (red arrows).

This contrasts with the immature biofilm that forms after 12 hr growth (**Figure 1**); vegetative cells are the dominate cell type in immature biofilms (vegetative cell and spore ratio of 700:1, respectively). This lack of spores within the biofilm, was reflected in the expression patterns of the early and late-stage sporulation genes. A number of early stage sporulation genes (*spo0A, soj, spoIIAA* and *spoIIAB*) are upregulated between 1.8 and 3.9-fold in biofilm cells, whilst the late stage genes (*spoIVB, spoIIIF, spoVG* and *obg*) were down regulated (between - 2.2 and -4.0-fold) compared with stationary phase cells^44^ (**Supplemental Table 3**). *C. difficile* sporulation and biofilm regulatory pathways intersect with Spo0A playing an important role in both pathways^45^, similar to biofilm formation in *B. subtilis*^25^. Spo0A is phosphorylated by orphan histidine kinases (KinA-D), thus low levels of Spo0A-P initiate biofilm formation whilst high levels initiate sporulation. CDR20291_1194 is an orphan histidine kinase with homology to the Kin proteins^44^ and was upregulated (2.2-fold) in biofilm cells; however, the potential interaction between Spo0A and CDR20291_1194 and the role in biofilm formation needs further investigation. In *B. subtilis*, Soj inhibits expression of Spo0A-P activated sporulation genes^46,47^, so it is interesting that during *C. difficile* biofilm formation, both *soj* and *spo0A* are upregulated, suggesting there could be a fine-tuning mechanism for biofilm initiation verse sporulation. Poquet *et al*.^14^ observed the induction of the late-stage sporulation cascade during biofilm maturation. During initial biofilm development, although cell aggregation was observed, there was a lack of self-produced matrix surrounding the cells and cells appeared intact, no cell debris, at this time.

Current evidence suggests that toxin production is increased within *in vitro* and *in vivo* formed biofilms^10,11,48,49^. In our experiments, both *tcdA* and *tcdB* were upregulated, 1.9- and 2.4-fold respectively, in biofilm cells (**Supplemental Table 3**). The remaining members within the PaLoc were also found to be upregulated in biofilm cells; however, significance was below our threshold (data not shown). Although highly expressed under both conditions, the CDT loci was not differentially expressed between biofilm and stationary phase cells (data not shown). Incorporation of *C. difficile* cells into mucosal biofilms would place these toxin producing cells adjacent to the host membrane, where small increases in toxin production could have an effect.

### Contribution of the putative biofilm regulator, CDR20291_2121 (sinR homologue), to biofilm formation

The *C. difficile* putative *sinR* homologue, CDR20291_2121, has previously been shown to interact with *sigD* (toxin production) and *spo0A* (sporulation) to control these pathways and the *sin* loci can modulate biofilm formation^14,28^. Transcriptome analysis of this locus showed decreased expression (−3.3-fold) of CDR20291_2121 in biofilm cultures compared with stationary phase cultures, although only low levels of expression were seen (**Figure 2**). This could explain the decreased levels of spores and increased toxin expression in biofilm samples. We sought to identify the specific role of CDR20291_2121, as an inhibitor of biofilm formation, by precision genome deletion. To prevent polar effects on downstream genes, we deleted 321/339 bp of CDR20291_2121, leaving the start and stop codons intact (**Figure 2 & Supplemental Figure 1**). Deletion of CDR20291_2121 significantly increased *C. difficile* biofilm formation compared to wild-type (*p* > 0.001), and plasmid complementation restored biofilm formation back to near wild-type levels. (**Figure 2**). SEM analysis of these biofilms, shows an increase in the amount of extracellular matrix, compared with WT (**Figure 1 and 2d**). Interestingly, plasmid-based over-expression of CDR20291_2121 in a wild-type background significantly reduced biofilm formation (*p* > 0.001), giving further evidence of the role of this gene in biofilm formation.

**Figure 2.**
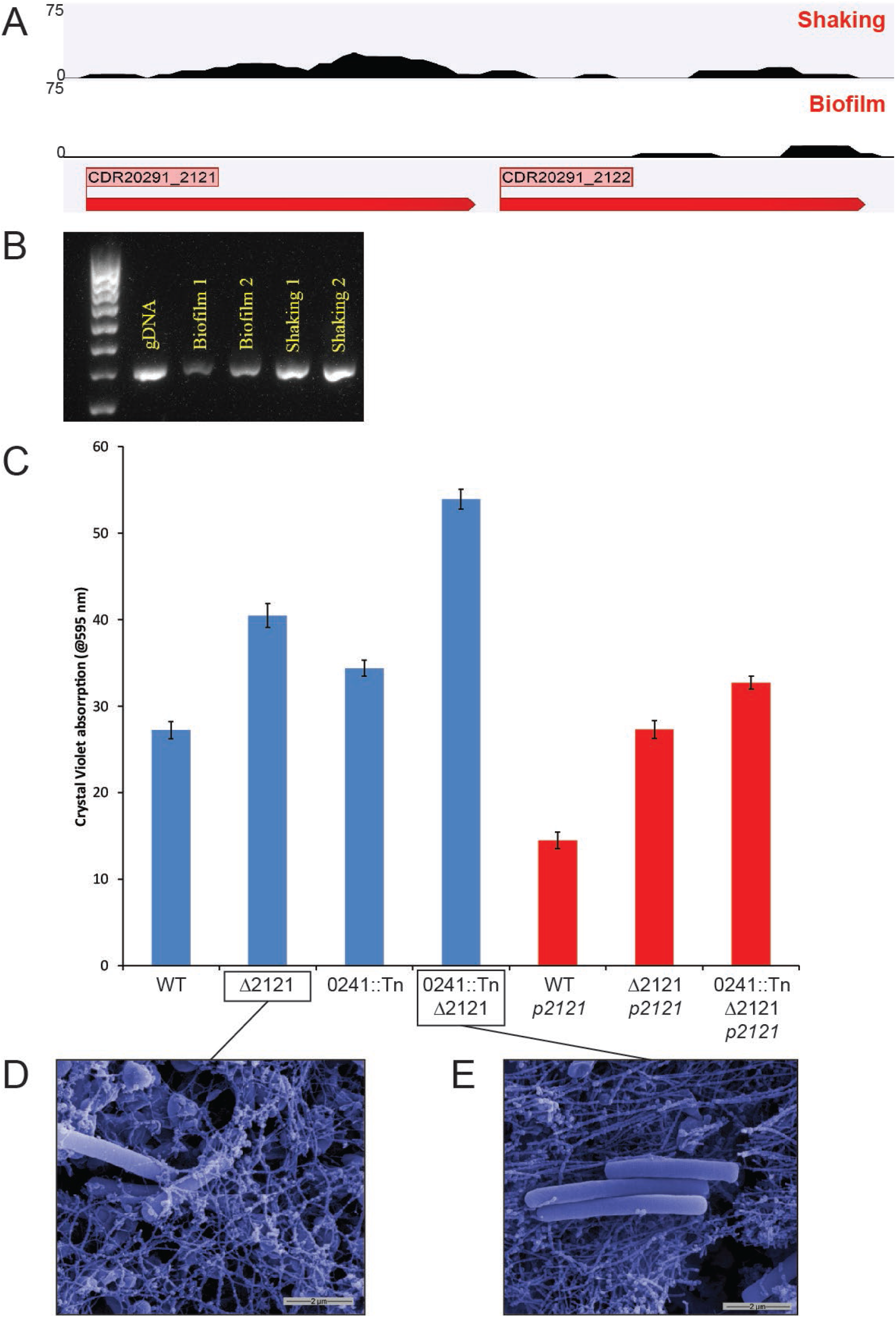
Characterising the role of SinR towards biofilm formation. Expression profiles of *sinR* (CDR20291_2121) and *sinR’* (CDR20291_2122) from RNA-Seq data (A) and independent RT-PCR reactions (B) (similar to **Figure 4**). Biofilm formation (C), as measured by crystal violet absorption, for *sinR* mutant (Δ2121 – blue bar) in either wild-type background (WT – blue bar) or a flagella glycosylation mutant background (0241::Tn – blue bars). *sinR* plasmid complemented (pRPF144-CDR20291_2121) in these background strains (red bars). *** indicates *p* < 0.001. False-coloured scanning electron microscopy images of biofilms from *sinR* mutant and *sinR*/flagella glycosylation double mutant strains (D).

The regulatory pathways that lead to biofilm formation in *B. subtilis* are multifactorial, whereby inhibition of flagella rotation represents a distinct entry mechanism (from de-repression of SinR) into biofilm formation^29^. Previously, we showed that prevention of flagella rotation, but not the complete absence of flagella, could lead to increased biofilm formation in *C. difficile*^17,18^. Deletion of CDR20291_2121 in this non-motile, flagella positive strain (CDR20291_0241::Tn) further enhanced biofilm formation, in a cumulative manner to produce a significantly greater biofilm than each individual mutation (**Figure 2**). Plasmid complementation restored biofilm formation similar to CDR20291_0241::Tn background strain. SEM images suggest this is due to further extracellular matrix production in this double mutant compared with the single Δ2121 mutant or wild-type (**Figure 1 and 2e**). These results show that, similar to *B. subtilis*, biofilm formation in *C. difficile* is a multifactorial process, whereby several environmental cues are needed for full biofilm formation.

### Changes to the metabolic landscape during initial biofilm formation

The entry into biofilm formation was associated with expressional changes to the metabolic pathways, compared with stationary phase cultures. Principal component analysis of our transcriptomic biofilm and stationary phase sequencing data showed variation between the biofilm and stationary phase culture data, suggesting that there were significant changes between these sample types (**Supplemental Figure S4**). From our expression data, 434 genes were differentially expressed between biofilm and stationary phase cultures; 205 genes were upregulated in biofilm cultures and 229 genes upregulated in stationary phase cultures (**Supplemental Table 3**). This number of differentially expressed genes (DEGs) between biofilm and stationary phase cultures is not surprising given that both culture conditions were grown in the same media for the same length of time and is similar to those reported in other studies^50^.

### Production of central metabolites

In comparison to stationary phase cultures, the metabolic landscape during the initial steps of biofilm formation is predominated by the upregulation of fermentation pathways that result in the production of central metabolites namely, acetyl-CoA, pyruvate and oxaloacetate. These core metabolites are used in a variety of metabolic pathways, such as the production of short chain fatty acids. During the early stages of biofilm formation, the metabolism of glycine was the predominant pathway for production of these central metabolites through three distinct pathways. Intracellular glycine can be converted to serine and then to pyruvate via the gene products of *glyA* and *tdcB*, which were upregulated, 2.1 and 2.6-fold respectively, in biofilm samples compared with stationary phase cultures (**Figure 3 & Supplemental Table 3**). Alongside this, glycine can be degraded to glyoxylate, which can enter the glyoxylate cycle, producing the central metabolite oxaloacetate; several putative genes encoding glyoxylate metabolising proteins were upregulated (2.1 – 3-fold) during biofilm formation (**Supplemental Table 3**). Like the citrate cycle, the glyoxylate cycle synthesises carbon-based macromolecules but from two-carbon sources instead, such as ethanol and acetate, thus bypassing the decarboxylation steps in the citrate cycle^51^. This glyoxylate shunt is important for the metabolic adaption of many pathogens during host-pathogen interactions^51,52^ and could be utilised by *C. difficile* during biofilm formation in the intestinal tract. Another route of glycine metabolism during biofilm growth utilises the combination of the glycine cleavage system (*gcvPBH*, upregulated 2.5 – 2.6-fold) and the wood-Ljundahl pathway (*hydN1, fdhF, fhs, fchA*, CDR20291_0648-0655; upregulated 2.1 – 3.1-fold). The glycine cleavage system produces the intermediate 5, 10-methylenetetrahydrofolate, which is utilised by the Wood-Ljundahl pathway to produce acetyl-CoA^53^. Glycine catabolism in this way appears to be important for biofilm formation, indeed mutational analysis of *gcv* loci in *B. subtilis* abolished biofilm formation in the presence of exogenus glycine^54^.

**Figure 3.**
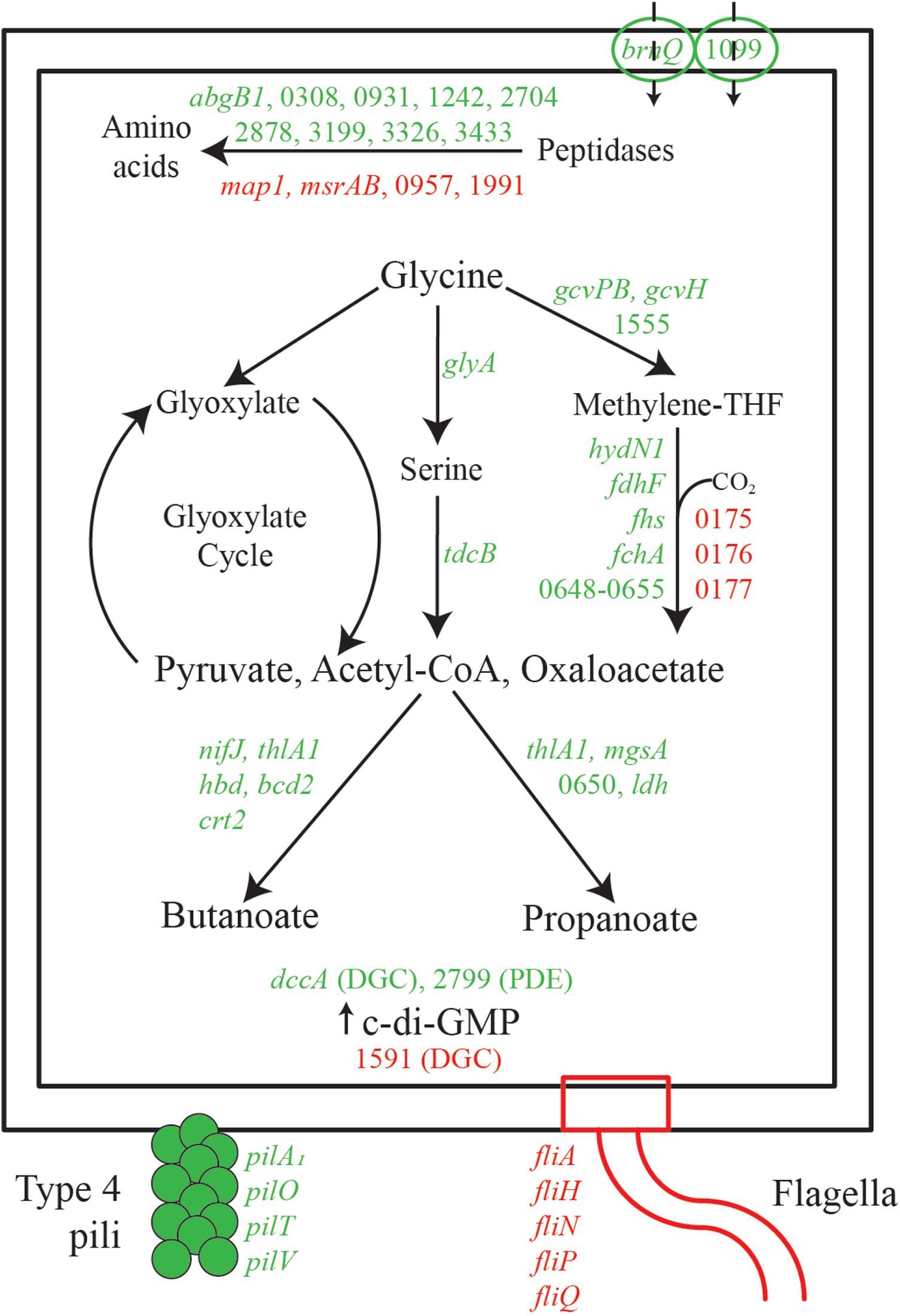
Dominant metabolic pathways and cell surface changes during biofilm formation. A model for amino acid uptake and metabolism of glycine, and cell surface organelle modifications during biofilm formation compared with stationary phase cells. Protein names/numbers are coloured green or red based on gene expression, up or down-regulated respectively, of biofilm cells compared with stationary phase cells. Protein names/numbers have been shortened (from CDR20291_*) for simplicity. THF – Tetrahydrofolate, CoA – co-enzyme A, c-di-GMP – cyclic diguanylate.

Interestingly, we detected the presence of a glycine riboswitch upstream of the CDR20291_1555-*gcvPB* operon (**Figure 4**). Glycine riboswitches respond to intracellular glycine levels and frequently occur upstream of, and control the expression of, *gcvTPB* operons in other bacteria. When glycine is bound, the riboswitch induces expression, causing the degradation of glycine^54,55^. The presence of CDR20291_CDs019 upstream of CDR20291_1555-*gcvPB* operon corresponded with increased expression of this operon in biofilm samples compared with shaking samples (**Figure 4**), suggesting intracellular accumulation of glycine during biofilm growth can occur. Structural predictions of this short glycine riboswitch showed homology to the Type I – singlet consensus structure with a ghost aptamer (antiterminator) structure at the 3’ end of the RNA (**Figure 4E**)^56^. High glycine levels correlated with a metabolic shift towards glycine metabolism in biofilm cells. In *Bacillus subtilis*, mutational analysis of this glycine riboswitch negatively affected swarming motility and biofilm formation in the presence of exogenous glycine^54^. A second glycine riboswitch was identified; CDR20291_CDs028 is found upstream of CDR20291_2174, a putative sodium:alanine symporter family, which was not expressed in either biofilm or shaking samples. Similarly, Khani *et al*.,^57^ identified a glycine riboswitch upstream of a sodium:alanine symporter family gene in *Strepococcus pyogenes*, which repressed this gene during high intracellular glycine levels. The size and predicted shape of this glycine riboswitch is characteristic of the two-tandem glycine-binding aptamers, followed by a single expression platform (**Figure 4F**)^55^.

**Figure 4.**
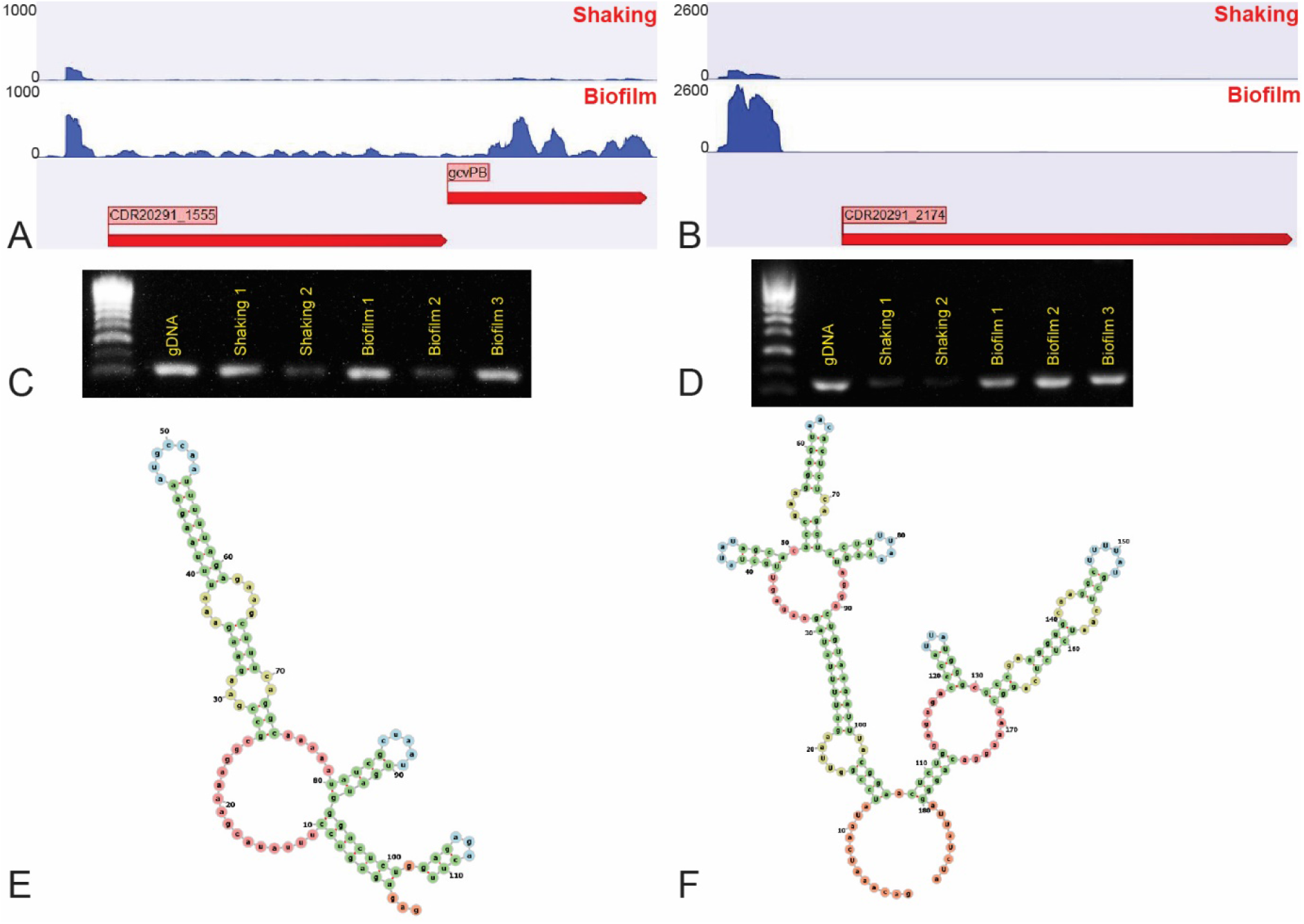
Expression and predicted structure of two glycine riboswitches. Expression and structure of glycine riboswitches CDR20291_CDs019 (A, C, E) and CDR20291_CDs028 (B, D, F). Expression patterns for each glycine riboswitch from RNA-Seq data (pictures are representative expression profiles from a single sequencing replicate), including the downstream gene expression (A, B) and RT-PCR for each sRNA (PCR picture shown is from independent RNA extractions to those used for RNA-Seq; replicate number is shown on the picture) with a genomic(g) DNA control (C, D). Predicted secondary structures of each riboswitch (using RNAfold software).

### Production of short-chained fatty acids

The production of pyruvate, acetyl-CoA and 2-oxobutanoate could be channelled into the metabolic pathways that produce short-chain fatty acids, namely the production of butanoate and/or propanoate (**Supplemental Table 3**). The conversion of pyruvate and acetyl-CoA to butanoate utilises several gene products (*nifJ, thlA1, hbd, bcd2* and *crt2*) in this pathway which are upregulated (1.9 – 2.7-fold) in biofilm cells compared with stationary phase cultures (**Figure 2**). Similarly, several genes in the early stages of the propanoate metabolism pathway which converts acetyl-CoA and 2-oxobutanoate to propanoate are upregulated (*ldh*, CDR20291_1007, *msgA, thlA1* and CDR20291_0650; 2.3 – 6.0-fold), whilst those genes involved in the late stages are highly expressed but were below our threshold in these experiments (data not shown). The capacity of *C. difficile* to utilise amino acids to produce fermentation products, such as butanoate and propanoate, when grown in minimal media has been reported^58^. However, these fermentation products appear to decrease as the biofilm matures, potentially indicating a reallocation of carbon resources in biofilm cells^14^.

### Reorganisation of the cell surface in sessile cells

Cells entering biofilm growth appear to reorganise their cell surface by altering the expression of cell wall proteins (cwps) and the major surface organelles, flagella and type 4 pili (T4P). A number of cwps are upregulated in biofilm cells, *cwp16, cwp20, cwp28, cwp84* and CDR20291_2099 & 2679 increased between 2.3 – 5.3-fold. Cwp16 is a putative N-acetylmuramoyl-L-alanine amidase and cwp20 is a putative penicillin-binding protein, both are highly conserved across *C. difficile* strains^59,60^. Cwp28 has an unknown function; structurally, Cwp28 possess the characteristic cell wall binding 2 motifs but these are not present in all *C. difficile* ribotypes^60^. Cwp84 is a cysteine protease involved in generation of the S-layer by cleaving the SlpA precursor; Cwp84 mutations exhibit decreased biofilm formation^12^, potentially due to a malformed S-layer. We have previously shown that the S-layer is important for protein retention at the cell surface^61^, indeed eight putative membrane proteins (between 2.2 and 7.1-fold) and four putative secreted proteins (between 2.6 and 3.4-fold) were upregulated in biofilm cells, including a haemolysin homologue, *hlyD*. Interestingly, five genes encoding membrane-associated proteins were down regulated in biofilm cells; CDR20291_0947 was down regulated by -7.2-fold compared with stationary phase cells (**Supplemental Table 3**).

Many genes encoding elements of the flagella system, including motor switch proteins, assembly proteins and flagella sigma factor, were inversely regulated in biofilm cells compared with stationary phase cells (*fliA, fliH, fliN, fliP, fliQ*; -1.6 to -3.3-fold) (**Figure 3**). The down regulation of the flagella operon correlates with the lack of flagella filaments on the cell surface, observed from SEM images (**Figure 1c**). Genes encoding elements of the T4P apparatus were upregulated (*pilA*_*1*_, *pilO, pilV, pilT, pilB, pilMN*; 2.4 – 5.1-fold) in biofilm cells (**Figure 3**). The switch from flagellate motile cells to non-flagellate twitching motility has been observed during *C. difficile* biofilm formation^16,62^.

The increased and decrease expression of the T4P and flagella operons, respectively, are controlled through c-di-GMP riboswitches; in our analysis, we detected the presence of five previously characterised c-di-GMP riboswitches^22,24,62,63^ (**Table 1**). The presence of the type I riboswitch, Cdi1_3, observed in the biofilm samples was associated with an absence of early stage flagella locus expression, giving further evidence this riboswitch acts as an ‘OFF’ switch during biofilm formation^22,24^. Concurrently, biofilm samples had increased expression of the type II riboswitch, Cdi2_4, which was associated with increased expression of several genes within the major pili operon. Interestingly, we detected expression of a second sRNA encoded upstream of Cdi2_4, with an unknown function (homologous to CD630_n01120/sCD4107) (**Supplemental Table 4**). Accumulation of intracellular c-di-GMP can initiate the early steps of *C. difficile* biofilm formation; we observed increased expression (28.2-fold) of *dccA*, which encodes a diguanylate cyclase to produce c-di-GMP^21^. Overexpression of *dccA* increases the intracellular concentration of c-di-GMP, which leads to decreased flagella synthesis and increased pili formation^63^. However, the upregulation of a putative phosphodiesterase, CDR20291_2799 (2.8-fold), which degrades c-di-GMP, suggests this intracellular signal is finely controlled; indeed, there are approximately 31 genes encoded on the *C. difficile* genome with putative roles in controlling c-di-GMP levels^21^.

**Table 2.**
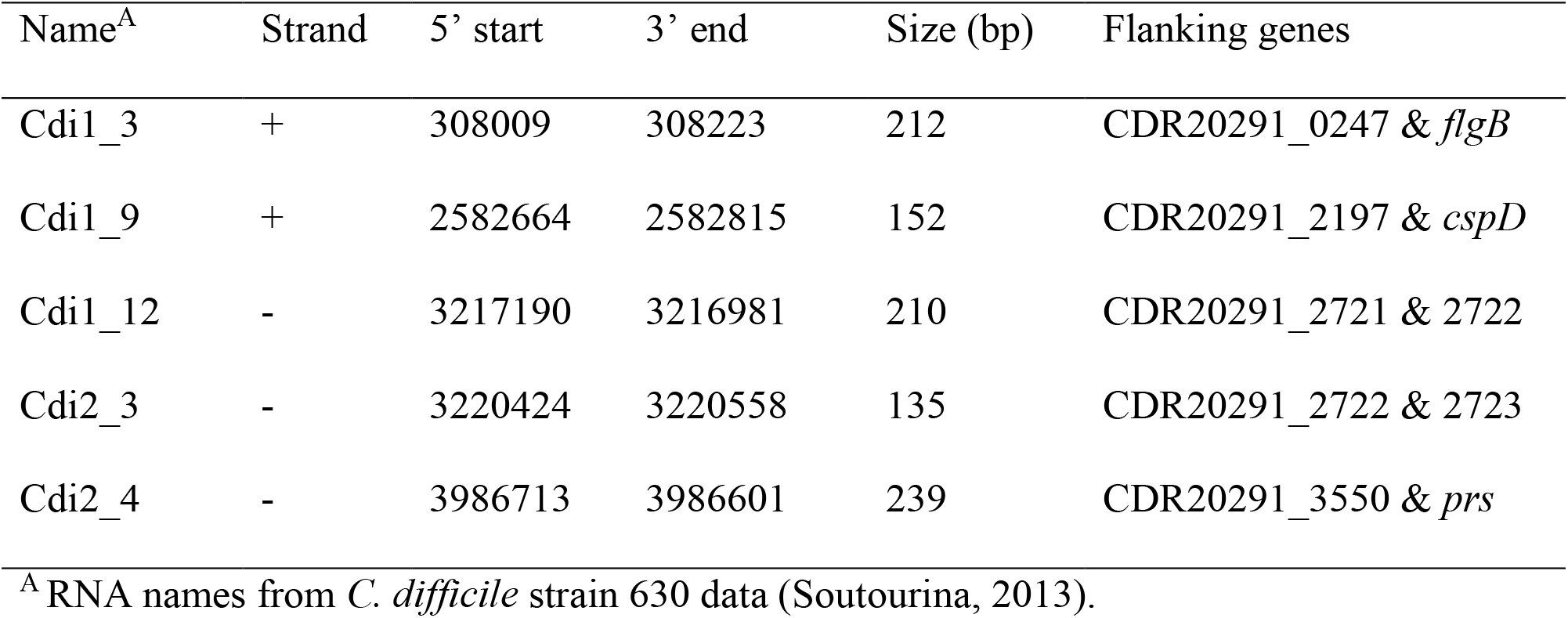
c-di-GMP riboswitches in *C. difficile* R20291

| Name<sup>A</sup> | Strand | 5' start | 3' end | Size (bp) | Flanking genes |
| --- | --- | --- | --- | --- | --- |
| Cdi1_3 | + | 308009 | 308223 | 212 | CDR20291_0247 & *flgB* |
| Cdi1_9 | + | 2582664 | 2582815 | 152 | CDR20291_2197 & *cspD* |
| Cdi1_12 | - | 3217190 | 3216981 | 210 | CDR20291_2721 & 2722 |
| Cdi2_3 | - | 3220424 | 3220558 | 135 | CDR20291_2722 & 2723 |
| Cdi2_4 | - | 3986713 | 3986601 | 239 | CDR20291_3550 & *prs* |

<sup>A</sup> RNA names from *C. difficile* strain 630 data (Soutourina, 2013).

## CONCLUSION

Here we describe the metabolic changes and regulatory aspects of a clinical *C. difficile* ribotype 027 epidemic strain during the transition from planktonic to biofilm cells. During biofilm initiation, cellular metabolic reprogramming, featuring an emphasis on glycine metabolism, was favoured to produce key central metabolites. Expression analysis shows that these metabolites are likely to be utilised for short-chain fatty acid production, butanoate and propanoate. These initial metabolic shifts are likely to undergo further changes as the biofilm matures. Indeed, Poquet *et al* (2018) reported an increase in fatty acid biosynthesis rather than fermentations in mature *C. difficile* biofilms. Secreted metabolic products could act as signals for biofilm formation, e.g. *B. subtilis* and *P. aeruginosa* respond to environmental acetic acid^64^ or pyruvate, respectively. Many bacterial species change their metabolic landscape during biofilm formation, due to oxygen and/or nutrient limitation, to promote cell survival (or death of a sub population)^65,66^. Inhibition of these metabolic cues could potentially be used to dismantle biofilms to enhance antimicrobial treatment. Factors like D-amino acids and cyclic-di-AMP are known to inhibit *B. subtilis* biofilm formation^67^; our data could provide the basis for similar interventions in respect of *C. difficile* biofilm formation, i.e. preventing c-di-GMP accumulation by inhibiting DccA.

Metabolic remodelling is largely controlled at the transcriptional level through changes to transcription factors and regulatory small RNAs^14,66,68^. Our data provide insights in to the sRNAs highly expressed in biofilm and stationary phase cells. Using glycine riboswitches, biofilm cells can control the intracellular levels of glycine for biosynthesis and energy whilst maintaining adequate amounts of the amino acid for protein synthesis. The different glycine riboswitches identified here are responsible for either inhibition or elevated expression of downstream genes depending on intracellular glycine concentrations. This could represent two distinct classes of riboswitches, like those for c-di-GMP, where genes can be differentially regulated (‘ON’ or ‘OFF’) depending on the intracellular concentration of a single signalling molecule. Some of the sRNAs could not be assigned a function based on homology. The discovery of new sRNA classes/motifs is constantly growing and those discovered in this study are likely to fall into this category. Weinberg *et al*.^69^ used a novel bioinformatic approach to identify 224 novel classes of non-coding RNA, and further research into these new classes is likely to reveal novel aspects of biological control. In our analysis, we identified 36 novel putative sRNAs depending on growth conditions. This suggests that *C. difficile* expresses subsets of different sRNAs during different cellular processes. It remains to be determined if any more sRNAs could be detected during other cell processes, such as germination or sporulation.

The metabolic reprogramming and cell surface remodelling seen during biofilm formation in *C. difficile* is regulated on multiple levels. The regulatory networks governing biofilm in *C. difficile* are unknown; however, there are similarities with biofilm regulatory pathways in *B. subtilis*. Our data suggest that the SinR homologue in *C. difficile* can repress biofilm formation. We have shown that this regulatory pathway can work independently of another regulatory mechanism, i.e. inhibition of flagella rotation^17^. Here, we highlight two regulatory pathways that can each contribute towards biofilm formation; however, fulminant biofilm formation occurs when both pathways are activated in *C. difficile*.

Our study describes the cellular metabolic and structural changes that occur during biofilm initiation, which supports earlier reports describing a connection between virulence attributes and metabolism^14,26,28,44,50^. Entry into biofilm formation is regulated at multiple stages and uses a combination of transcriptional regulators and small non-coding RNAs to pass these check points. By understanding the mechanisms behind this, we hope to find potential interventions to dysregulate biofilm formation, possibly affecting multiple virulence pathways, given the inter-connected nature of these processes.

## ACKNOWLEDGEMENTS

We acknowledge the help of M. Mullin (Integrated Microscopy, University of Glasgow) for SEM sample preparation. The authors gratefully acknowledge the support of the Wellcome Trust (grant number 086418 awarded to G.R.D) and the Rosetrees Trust (grant number M636 awarded to M.H.W.).

