## Supplementary material for "Insights into the regulatory mechanisms of *Clostridioides difficile* biofilm formation"

*Table S1 – strains and plasmids used in this study*

| Strain/Plasmid | | Characteristics | Reference/Source |
| --- | --- | --- | --- |
| **Strains** | | | |
| *E. coli* | | | |
|  | TOP10 | F¯ *mcrA* ∆(*mrr-hsdRMS-mcrBC*) φ80*lacZ*∆M15 ∆*lacX74 nupG recA1 araD139 ∆*(*ara-leu*)*7697 galE15 galK16 rpsL*(Str^r^) *endA1* λ¯ | Invitrogen |
|  | CA434 | *E. coli* HB101 [F¯ *mcrB mrr hsdS20*(r_B­­_¯m_B_¯) *recA13 leuB6 ara-14 proA2 lacY1 galK xyl-5 mtl-1 rpsL20*(Sm^r^) *glnV44* λ¯] | Faulds-Pain 2013 |
| *C. difficile* | | | |
|  | R20291 | Wild-type; PCR ribotype 027 (Stoke Mandeville, U.K., epidemic strain) |  |
|  | R20291::Tn 0241 | R20291 mutant, containing a ClosTron insertion in flagella glycosylation gene (CDR_*0241*) | Valiente 2016 |
|  | R20291 Δ2121 | R20291 mutant containing a 321 bp deletion in biofilm master regulator gene (CDR_*2121*) | This study |
|  | R20291::Tn 0241 Δ2121 | R20291 double mutant containing a ClosTron insertion in CDR_*0241* and a 321 bp deletion in CDR_*2121* | This study |
|  | R20291 pCDR_2121 | R20291 WT harbouring pRPF144-CDR_*2121* | This study |
|  | R20291 Δ2121 pCDR_2121 | R20291 Δ2121 harbouring pRPF144-CDR_*2121* | This study |
|  | R20291::Tn 0241 Δ2121 pCDR_2121 | R20291::Tn 0241 Δ2121 harbouring pRPF144-CDR_*2121* | This study |
| **Plasmids** | | | |
|  | pRPF144-*gusA* | Constitutive expression vector for *C. difficile* expressing *gusA* | Fagan 2011 |
|  | pRPF144-*ftsZ-LOV* | pRPF144 with *gusA* replaced with *C. difficile* R20291 *CDR_2121* | This study |
|  | pMTL82151 | *E. coli* – *C. difficile* shuttle plasmid (pCD6; *catP*; ColE1+*tra*) | Faulds-Pain 2013 |
|  | pMTL82151-CDR_2121 | pMTL82151 with ± CDR_2121 1200 bp | This study |

*Table S2 – primers used in this study*

| Primer name | Characteristics | Sequence (5’-3’) |
| --- | --- | --- |
| **Mutant construction** | | |
| CDR2121 HA1 F | Primer set to amplify 1200 bp upstream and start of CDR_2121 – homology arm 1 | TGATTACGAATTCGAGCTCGGTACCCCCAGTTTCAGGTGATATAA |
| CDR2121 HA1 R |  | CTCACTTATATTTCTTTGTATATATTTGCCAATTATTATCCCTCC |
| CDR2121 HA2 F | Primer set to amplify stop codon of CDR_2121 and 1200 bp downstream – homology arm 2 | GATAATAATTGGCAAATATATACAAAGAAATATAAGTGAGGGAAA |
| CDR2121 HA2 R |  | GCGTGACGTCGACTCTAGAGGATCCGGTTATATATGTAATATAAT |
| LSHTM F | Primer set to confirm homology arm insertion into plasmid pMLT82151 | CAGGAAACAGCTATGACC |
| LSHTM R |  | TGTAAAACGACGGCCAGT |
| CDR2121 KO check F | Primer set to confirm chromosomal mutation and sequence | TACAACAACGTCTAACTTCC |
| CDR2121 KO check R |  | GCCATTAAAACCTACTAGAC |
| CDR2121 KO check R2 |  | AAATATGGTTGGAGTATCGC |
| **Plasmid complementation** | | |
| CDR2121 p144 exp F | Primer set used to amplify CDR_2121 fragment for expression cloning | GGAAAAAATAATAAGAGCTCATAATCTAAAGTGGAGGGATAATAA |
| CDR2121 p144 exp R |  | CGGCCGTTACTAGTGGATCCTTATATTTCTTTGTATTTAATGATG |
| NF793 | Primer set to confirm insertion of fragment into plasmid pRPF144 | CACCTCCTTTTTGACTTTAAGCCTACGAATACC |
| NF794 |  | CACCGACGAGCAAGGCAAGACCG |
| **RT-PCR confirmation of sRNAs** | | |
| CDR20291_CDs001 forward | Amplification of CDR20291_CDs001 located between CDR20291_0132 and 0133 | TAGGTCTGTGGTTGAAAG |
| CDR20291_CDs001 reverse |  | CATAAAGGTATGCTAGCT |
| CDR20291_CDs005 forward | Amplification of CDR20291_CDs005 located between CDR20291_0405 and 0406 | CATTTTTGAGGTTGTCGT |
| CDR20291_CDs005 reverse |  | CAGGTAAGCTCCGAATAG |
| CDR20291_CDs012 forward | Amplification of CDR20291_CDs012 located between CDR20291_tlpB and 0810 | AAGAACCGTAGGAACTAC |
| CDR20291_CDs012 reverse |  | ACAAAGTAGTCTGAACCT |
| CDR20291_CDs017 forward | Amplification of CDR20291_CDs017 located between CDR20291_1445 and 1446 | CAGTGAGCGATATTTGTG |
| CDR20291_CDs017 reverse |  | TGCAGTGAACCATGAGTA |
| CDR20291_CDs019 forward | Amplification of CDR20291_CDs019 located between CDR20291_1554 and 1555 | GAGAGAGTCCTTTATACG |
| CDR20291_CDs019 reverse |  | TCTCCAGAGTCCCATCAA |
| CDR20291_CDs020 forward | Amplification of CDR20291_CDs020 located between CDR20291_1564 and 1565 | AAGACTTGATGTAAGCAG |
| CDR20291_CDs020 reverse |  | GGGGATGGATAATATGTT |
| CDR20291_CDs026 forward | Amplification of CDR20291_CDs026 located between CDR20291_1978 and 1979 | TAGGTCTGTGGTTGAAAG |
| CDR20291_CDs026 reverse |  | CATAAAGGTATGCTAGCT |
| CDR20291_CDs028 forward | Amplification of CDR20291_CDs028 located between CDR20291_2173 and 2174 | GAAGAGTTGCTATATAGC |
| CDR20291_CDs028 reverse |  | GATAATCCCTGTCCTTTT |
| CDR20291_CDs030 forward | Amplification of CDR20291_CDs030 located between CDR20291_2492 and trpS | ACAATGTATGTTAGAAGT |
| CDR20291_CDs030 reverse |  | AAAAAAGATGAGCTTTTG |
| CDR20291_CDs032 forward | Amplification of CDR20291_CDs032 located between CDR20291_2501 and ileS | TTTTTGAGGTTGTCGTTC |
| CDR20291_CDs032 reverse |  | AACTTCAACGACTATTCC |
| CDR20291_CDs033 forward | Amplification of CDR20291_CDs033 located between CDR20291_2688 and 2689 | ACAAACAACTAAATCAAT |
| CDR20291_CDs033 reverse |  | AACAATTAGAAATGGAGG |
| CDR20291_CDs034 forward | Amplification of CDR20291_CDs034 located between CDR20291_cobT and 3262 | CCTATTTTGAAGTCACAA |
| CDR20291_CDs034 reverse |  | TAATGTTAAATGGGAAGT |
| CDR_luxS-3437 forward | Amplification of sRNA located between CDR20291_luxS and 3437 | ATTATTTCAAGTCATCTC |
| CDR_luxS-3437 reverse |  | AAGTCAGTTTAATAAGAT |

*Supplementary results*

Other transcriptomic changes occurring during initial biofilm formation

**Amino acid metabolism and transport**

Stationary phase cultures appear to utilise Stickland reactions to metabolise glycine for energy production (Bouillaut 2013, Poquet 2018, Hoffmann 2018). Indeed, the genes involved in glycine reduction via the Stickland pathway were all inversely regulated during biofilm formation (Supplemental Table 3). Aside from glycine, proline can be used as an electron acceptor in Stickland reactions, whereby proline reduction requires the proline reductase enzyme complex (*prd* loci). The induction of the *prd* loci under biofilm conditions has previously been noted in strain 630 (Poquet 2018). In this study, the entire *prd* loci is upregulated biofilm cells, although only *prdE/F* were above the 2-fold change threshold. This suggests that proline reduction is an additional source of energy favoured during biofilm formation, whereas during stationary phase, glycine reduction is favoured (Hoffmann 2018).

The exogenous L-cysteine present in our growth media is utilised during biofilm growth, as shown by the induction of L-cysteine metabolic pathways and sulphur-containing uptake systems. *ssuA* and CDR20291_3096 encode a sulfonate-binding transport system and a putative permease, which are upregulated 2.2 and 2.4-fold, respectively, in biofilm cells compared with stationary phase cells. In biofilm cells, intracellular cysteine is transaminated to form pyruvate directly by CDR20291_3263 (21.9-fold), or via the intermediate 3-mercaptopyruvate by CDR20291_2719 (2.2-fold) and *sseA* (2.2-fold) to form pyruvate (Supplemental Table 3). Other amino acids metabolised to produce core metabolites include threonine and aspartate. CDR20291_2719 is upregulated (2.2-fold) in biofilms cells and encodes a putative aspartate aminotransferase that coverts L-aspartate to oxaloacetate, which can be used in the glyoxylate cycle. *tdcB* encodes a threonine dehydratase that coverts threonine into 2-oxobutanoate, a metabolite used for the production of propanoate, and was also upregulated in biofilm cells (2.6-fold).

In order to provide the cell with these amino acids, a number of peptide/amino acid transporters and peptidases were upregulated in biofilm cells. This is not unexpected given that the media used in our assay contains tryptone, which is rich in peptides. Two putative branched-chain amino acid transport carrier genes (CDR20291_1099 & *brnQ*) were upregulated in biofilm cells by 2.8 and 3.3-fold, respectively, along with a putative amino acid permease (CDR20291_2931; 3.1-fold) and an inversely regulated alanine or glycine:cation symporter system (CDR20291_1594; -2.3-fold). *C. difficile* strain R20291 harbours several DEGs encoding peptidases; 8 were upregulated (between 2.1 and 5.6-fold), and 4 inversely regulated (between -2.5 and -3.0-fold) (Supplemental Table 3). Interestingly, those peptidases that are inversely regulated have putative aminopeptidase functions preferentially for the release of methionine; however, within biofilm cells methionine is produced from the conversion of L-homocysteine through the SAM recycling pathway via the upregulation of *luxS* (3.5-fold) and CDR20291_3434 (3.0-fold) (Supplemental Table 3).

**Energy production**

Energy production during initial biofilm formation is generated by a bifunctional V-type ATP synthase encoded by the ntp loci, potentially using a sodium gradient to generate ATP. All eight genes (*ntpA-G)* in this operon were upregulated in biofilm cells (2.3 – 3.3-fold) compared with stationary phase growth. The V-type ATP synthase is also found in Enterococcus hirae and Thermus thermophilus (Murata 2005, Hosaka 2006).

**Translation**

During biofilm formation, many genes encoding the small and large ribosomal subunits, including the translation initiation factors *infA* and *tsf*, were inversely regulated between -2.0 and -6.4-fold, compared with stationary phase cells. In addition, we observed the down regulation of several major chaperone-encoding genes (*clpB/C, groEL, grpE dnaK, dnaJ*; between -1.9 and -2.6-fold) in biofilm cells.

**Cell wall and membrane biogenesis**

The metabolic pathways discussed so far are consistent with a non-growing phenotype, as discussed by Hoffmann et al. (2018). Augmenting this phenotype is the down regulation of lipid biogenesis genes; all fab genes were inversely regulated in biofilm cells, with *fabG* and *fabH* significantly inversely regulated, -2.8- and -2.1-fold (Supplemental Table 3). Interestingly, the acyl carrier protein, *acpP*, and a putative lipid transport protein similar to DegV in *B. subtilis* (CDR20291_2497), were upregulated in biofilm cells, 3.1- and 3.0-fold respectively. During biofilm maturation, the cell wall and envelope of biofilm cells undergoes remodelling compared with planktonic cells (Poquet 2018); the upregulation of *acpP* and CDR20291_2497 could be involved in initiating this process, potentially during cell differentiation observed in *C. difficile* biofilms (Pedrido 2013, Jahn 2003) rather than cell replication. In accordance with this, those genes involved with synthesis of peptidoglycan are also inversely regulated in biofilm cells; genes involved in amino-sugar metabolism and peptidoglycan synthesis were down regulated (*nagA*, *nan* operon and *glmS/D*; between -3.7 and -2.3-fold). A mechanism for potential membrane remodelling is the incorporation of D-alanyl ester-linked teichoic acids into the cell wall, which decreases the negative charge on the bacterial cell surface. In Gram-positive bacteria, this is achieved through the expression of the *dlt* operon; the *C. difficile dlt* operon was upregulated in biofilm cells, although only *dltB* significantly so (2.7-fold), along with a putative D-ala-D-ala carboxypeptidase (3.3-fold). Similar to other biofilm-forming bacteria, this change in cell surface charge could contribute towards *in vivo* biofilm resistance to the cationic antimicrobial peptides that are found in the mammalian intestinal tract (Fabretti 2006, Nilsson 2016). Additionally, *dltB* expression has been suggested to help regulate biofilm formation in Staphylococcus aureus (Huang 2014).

**Transporters**

A number of transporting systems are differentially regulated during biofilm formation. During biofilm formation, two sugar transport system were upregulated; the entire ribose ABC transporter system (*rbsA-C*, 3.9 – 6.8-fold; and the regulator *rbsR*, 2.2-fold) and a N-acetylglucosamine PTS system (CDR20291_2927-2928, 2.4-fold each). Poquet et al. (2018) also observed an upregulation of this PTS transport system; however, they observed the down regulation of the *rbs* loci during biofilm maturation. This may be due to differences in the stages of biofilm development investigated or between the strains used in the respective studies. Those genes encoding iron uptake systems are differentially regulated between growth phases; in the initial stages of biofilm growth, genes encoding iron transport systems, such as *fhu* operon, *feoB1/A3* and CDR20291_1326/1639, were inversely regulated compared with stationary phase cells (-2.2 – -11.8-fold). The down regulation of iron transport systems during biofilm initiation suggests iron acquisition is not a priority, which is supported by the upregulation of ferredoxin-dependent amino acid fermentations (*etf* genes, *hbd, bcd2, nifJ,* CDR20291_0114) (Berges 2018). During biofilm formation, 11 genes encoding ABC transporter systems were differentially regulated, including a single upregulated putative efflux ABC transporter system (CDR20291_1376-1378; 2.7 – 3.2-fold), with the remaining genes inversely regulated (-2.2 – -6.2-fold). Interestingly, a sulfonate ABC transport system was down regulated (CDR202091_2825-2826; -2.9 – -4.1-fold). Other DEGs in biofilm cells belong to the symporter family of transport systems; CDR20291_2109, a putative sodium:solute symporter, showed increased expression in biofilm cells (2.8-fold), whereas a further five symporter systems had reduced expression (-2.6 – -8.6) compared with stationary phase cells.

**Transcriptional regulators**

The metabolic pathways and cellular changes induced during the switch towards biofilm formation must be tightly regulated and can involve multiple components and signalling molecules. In our experiment, we identified 15 upregulated and 10 down regulated transcriptional regulators belonging to either the two-component response systems or the helix-turn-helix (HTH) motif regulators. CDR20291_1748-1749 (a two component regulator with unknown function), were the most down regulated of all regulator genes (-4.6 and -36.6-fold, respectively) and are not present on the *C. difficile* strain 630 genome; whereas 12 putative HTH regulators, which bind metabolites to activate/repress gene expression, were upregulated (2.1 – 11.5-fold) in biofilm cells (Supplemental Table 3). CDR20291_3046, encoding a *MerR*-family HTH regulator, was upregulated 11.5-fold in biofilm cells. This family of HTH regulators responds to a variety of signals, such as heavy metal or nutrient stress (Brown 2003), but have been implicated in biofilm formation (Huang 2012). Another HTH transcriptional regulator belonging to the CRP/FNR family, CDR20291_0552, was highly upregulated (6.2-fold) in biofilm cells. This number of differentially regulated genes encoding transcriptional regulators potentially signifies the level of molecular control a cell uses to commit to a lifestyle change, such as biofilm formation, and may even require multiple signals for full biofilm formation to occur. Such levels of control are also seen during sporulation (Saujet 2013, Pettit 2014) and in *B. subtilis* biofilm formation (Cairns 2014).

**Mobile genetic elements**

*C. difficile* strain R20291 harbours two large prophage islands (CDR20291_1201-1225 & CDR20291_1415-1462), which nearly all genes were inversely regulated in biofilm cells (-2.0 – -6.1-fold). These prophage islands are proposed to contribute to the extracellular DNA in the biofilm matrix (Dawson 2012, Slater 2019); the down regulation of these islands during biofilm initiation might be a consequence of the early time point investigated here, and the expression of these islands may be induced during biofilm maturation. Also of note is the increased expression of a putative conjugation island (CDR20291_1797-1799; 2.2 – 2.8-fold) in biofilm cells.

Identification of non-coding small RNAs expressed during biofilm and stationary phase growth

Deep sequencing of our cDNA libraries was used to detect small non-coding RNAs that are potentially involved in regulation of biofilm formation. Visual analysis of RNA-Seq reads aligned along the R20291 genome revealed multiple alignments outside the coding sequences. A total of 107 putative sRNAs were identified in these experiments; 10 of these were found to be upregulated in stationary phase cultures, 25 upregulated during biofilm growth and 72 expressed under both growth conditions (**Supplemental figure 4** & **Supplemental Table 4**). The large number of sRNAs expressed under both of the growth conditions tested is not surprising given our cultures were grown in the same growth media for the same length of time. Of the 107 identified sRNAs in strain R20291, 36 sequences represent novel sRNAs with either no homology in strain 630 or with homology but without reported identification, whilst the remainder (71) show homology to those reported previously in strain 630 (Chen 2011, Soutourina 2013). For clarity, we used the same sRNA nomenclature originally used for strain 630, except for the novel sRNAs, which have been given a strain/sRNA identifier.

**Common sRNAs expressed during both growth conditions**

The detection of 72 common sRNAs during both growth conditions suggests these could form a ‘core’ RNome needed for *C. difficile* R20291 metabolic/growth transcriptional regulation, under the conditions examined here. These shared sRNAs include 16 putative novel sequences, 8 of which have no homologous sequences in strain 630 (Chen 2011 & Soutourina 2013) (**Supplemental Table 4**). Interestingly, we detected a previously uncharacterised sRNA within the prophage 1 island (CDR20291_CDs018), containing an *abiF* motif (Weinberg 2017). Prediction of the target mRNA revealed CDR20291_1460 mRNA (z-score -3.35; uncharacterised protein) as a potential target, albeit with low z-score. Most of these common regulatory transcripts appear to be *cis*-acting, reside upstream of genes with a known metabolic function and are homologous to several sRNA motifs, i.e. T-box leader sequences and several riboswitches (lysine, SAM, cobalamin and purine). There is a high representation of sRNAs with a putative T-box leader sequence. This motif regulates the expression of aminoacyl-tRNA synthetase genes and controls gene expression of amino acid uptake and biosynthesis (Green 2013). In our experiments, we detected the differential expression of several tRNA synthetase genes, which could be controlled by these sRNAs, potentially causing the different amino acid metabolic pathways observed between the two growth conditions.

**Identification of biofilm specific small RNAs**

RNA motif homology searches among the biofilm specific sRNA sequences (Kalvari 2017), yielded putative matches for CDR20291_CDs001, CDs013 and CDs016. CDR20291_CDs001 displayed homology to the *raiA* RNA motif family; expression of this sRNA was confirmed by RT-PCR in biofilm samples and, to a much lesser extent, shaking. This newly discovered RNA motif is present in two phyla (Firmicutes and Actinobacteria) and possess structural characteristics similar to those sRNAs with sophisticated functions; however, its mode of action and potential targets are unknown (Weinberg 2017). CDR20291_CDs001 resides upstream of a putative transcriptional antiterminator (CDR20291_0133), but this gene was not differentially regulated in either of our growth conditions and mRNA target predictions yielded multiple mRNA targets at other genetic loci (all with a low z-score). Whilst we confirmed the expression of this sRNA, we could not determine whether CDR20291_CDs001 is likely to act as a *cis* or *trans*-regulatory sRNA or deduce its likely target. CDR20291_CDs013 and CDs016 reside upstream of a putative transposase (CDR20291_0959) and a pseudogene (CDR20291_1338), respectively, and contains a group I intron. These self-splicing ribozymes are found widely inserted into Gram-positive bacteria, but no biological function has been identified to date (Henrik 2009). A further 3 sRNAs were located adjacent to putative transposases genes (**Supplemental** **Table 4**). sRNAs within transposable elements can act as silencers to prevent transposition, thus mitigating the genome instability caused by uncontrolled transposition; however, during stress conditions, sRNAs can act in *cis* and *trans* to regulate host genes (Wheeler, 2013). The characterisation of these regulatory elements, and the effects on the *C. difficile* transcriptome, warrants further investigation.

Predictions of the likely targets of the remaining novel sRNAs, identified three sRNAs that are predicted to regulate downstream genes in a *cis* manner (**Supplemental Table** **4**). CDR20291_CDs026 and CDR20291_CDs018 lie upstream of a putative selenium-dependent hydroxylase accessory gene (*yqeC*; z-score -11.5) and a putative cation efflux gene (a low z-score of -2.05), respectively, and are likely to have a negative effect on the downstream expressed mRNA, as no expression at these loci were observed. A further *cis* regulating sRNA, CDR20291_CDs027, resides upstream of a putative copper transporting P type ATPase (z-score -7.68), appearing to negatively affect this gene under biofilm formation (-1.51-fold, not significant).

Although not a novel sRNA to this study, we detected the expression of a sRNA upstream of the *luxS* operon, similar to that detected in a previous study on *C. difficile* strain 630 (Soutourina 2013). An increase in this sRNA was associated with increased expression of *luxS* transcript in our biofilm samples compared with the stationary phase shaking samples. LuxS catalyses the formation of an AI-2 autoinducer molecule, which is produced, and detected, by many different bacterial species to coordinate, on a population level, a variety of cellular processes, including biofilm formation. The production of AI-2 in *C. difficile* is growth dependent (Carter 2005), and a functional LuxS is required for full biofilm formation (Dapa 2013, Slater 2019).

*Figure S1*

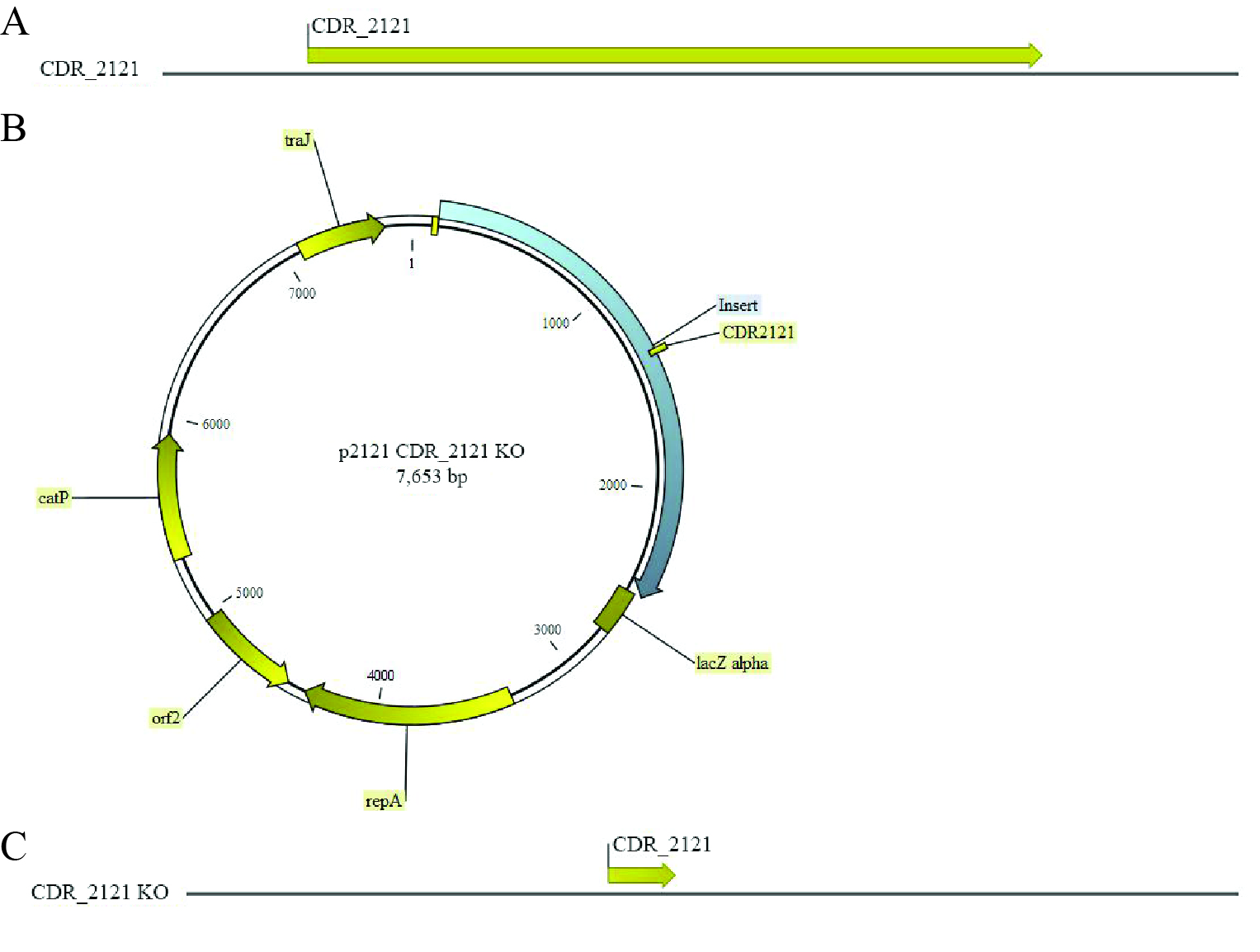

**Figure S1.** *In silico* schematic of the CDR20291_2121 wild-type genetic locus (**A**), the homology recombination plasmid with the deleted sequence (**B**) and the resulting chromosomal mutation (**C**).

*Figure S2*

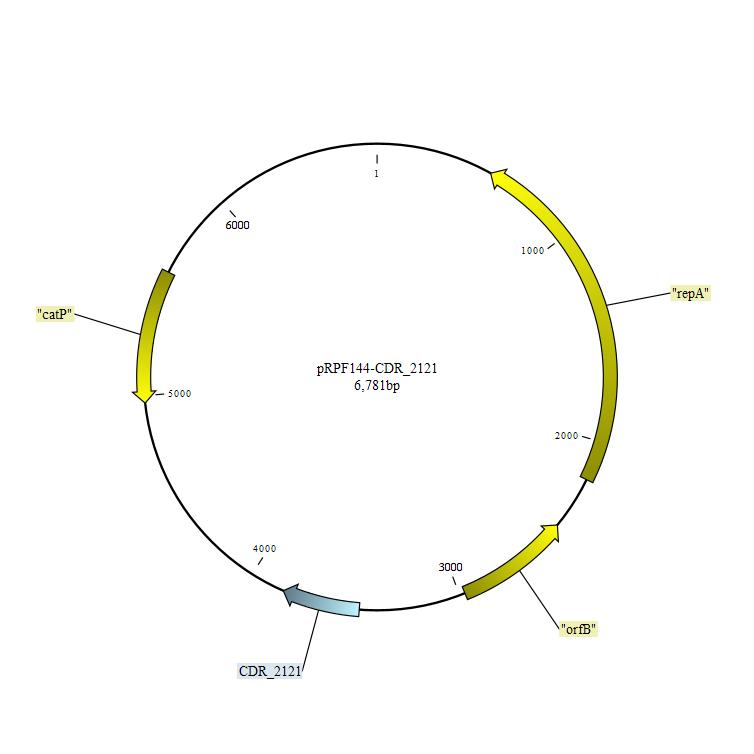

**Figure S2.** *In silico* schematic of the pRPF144-CDR_2121 expression plasmid used for complementation studies.

*Figure S3*

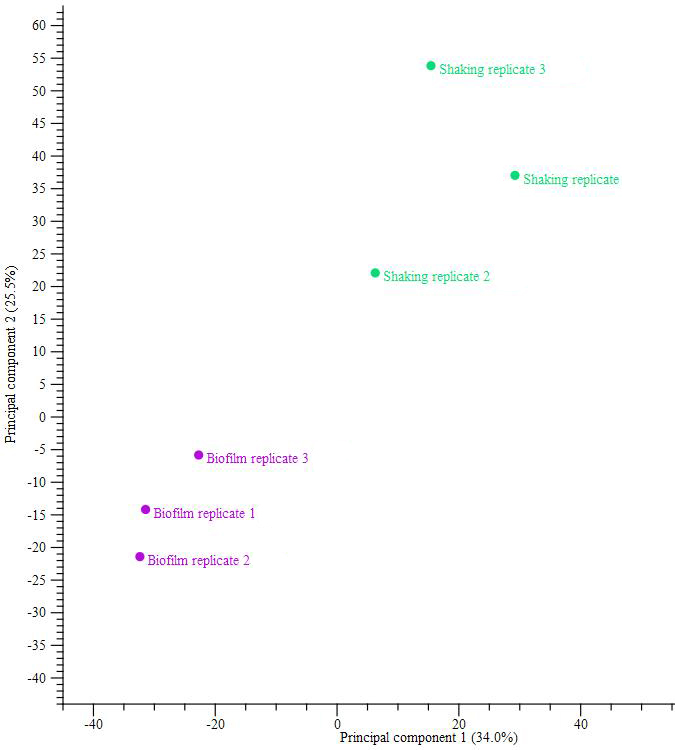

**Figure S3**. Principal component analysis based on the RNAseq data from biofilm (blue dots) and stationary cultures (green dots).

*Figure S4*

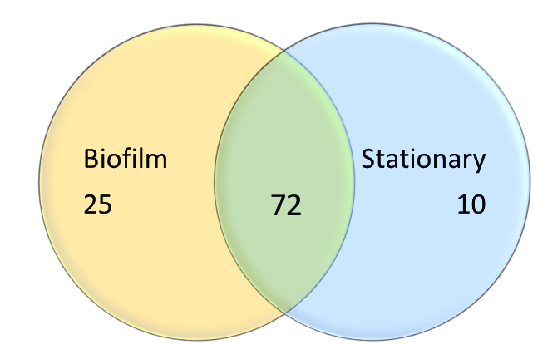

*Table S3 – Excel file ‘Supplementary Table 3’*

All differentially expressed genes and associated pathways.

*Table S4 – Excel file ‘Supplementary Table 4’*

All small RNAs detected in this study.
