## Supplemental Table 3 for "Insights into the regulatory mechanisms of *Clostridioides difficile* biofilm formation"

| Gene ID[1] | Name | Gene product/function | Differential expression by RNAseq (fold change) |
|---|---|---|---|
| **Central Carbon Metabolism** | | | |
| **Starch and sucrose metabolism** | | | |
| 2415 | gbeA | 1,4-alpha-glucan branching enzyme | 2.84 |
| 2930 | treA | trehalose-6-phosphate hydrolase | 2.65 |
| 2401 | | maltose-6'-phosphate glucosidase | -7.45 |
| **Glycolysis/Gluconeogenesis** | | | |
| 2570 | nifJ | pyruvate-ferredoxin/flavodoxin oxidoreductase | 2.25 |
| 0762 | aksA | homocitrate synthase | -3.76 |
| 3027 | gpmI | 2,3-bisphosphoglycerate-independent phosphoglycerate mutase | -2.06 |
| 1662 | gapA | glyceraldehyde 3-phosphate dehydrogenase | -2.42 |
| 3030 | gapB | glyceraldehyde 3-phosphate dehydrogenase | -1.92 |
| 0507 | gapN | glyceraldehyde-3-phosphate dehydrogenase (NADP+) | -2.96 |
| 2207 | | putative phosphoglucomutase | -2.24 |
| **Glyoxylate and dicarboxylate metabolism** | | | |
| 0114 | | 2-oxoglutarate ferredoxin oxidoreductase subunit delta | 2.06 |
| 0847 | | alanine--glyoxylate aminotransferase family protein | 2.12 |
| 1010 | | glycolate oxidase | 2.39 |
| 1867 | | glyoxalase | 2.96 |
| 3443 | | phosphoglycolate phosphatase | -3.12 |
| 3449 | | putative glyoxalase | -3.14 |
| **Wood-Ljundahl pathway** | | | |
| 3174 | hydN1 | electron transport protein HydN | 2.53 |
| 3179 | fdhF | formate dehydrogenase (NADP+) | 2.36 |
| 0645 | fhs | formate--tetrahydrofolate ligase | 2.50 |
| 0646 | fchA | methenyltetrahydrofolate cyclohydrolase | 2.36 |
| 0648 | | 5,10-methylenetetrahydrofolate reductase | 2.06 |
| 0649 | | 5,10-methylenetetrahydrofolate reductase | 2.09 |
| 0650 | | dihydrolipoamide dehydrogenase | 2.29 |
| 0651 | | CO dehydrogenase maturation factor | 2.54 |
| 0652 | | acetyl-CoA decarbonylase/synthase, CODH/ACS complex subunit delta | 2.56 |
| 0653 | | acetyl-CoA decarbonylase/synthase, CODH/ACS complex subunit gamma | 2.93 |
| 0654 | | 5-methyltetrahydrofolate corrinoid/iron sulfur protein methyltransferase | 2.87 |
| 0655 | | acetyl-CoA synthase | 3.10 |
| **Butanoate metabolism** | | | |
| 0915 | thlA1 | acetyl-CoA C-acetyltransferase | 2.25 |
| 0914 | hbd | 3-hydroxybutyryl-CoA dehydrogenase | 2.16 |
| 0913 | crt2 | enoyl-CoA hydratase | 1.87 |
| 0910 | bcd2 | butyryl-CoA dehydrogenase | 2.66 |
| 0175 | | anaerobic carbon-monoxide dehydrogenase catalytic subunit | -3.15 |
| 0176 | | anaerobic carbon-monoxide dehydrogenase iron sulfur subunit | -2.48 |
| 0177 | | putative oxidoreductase, NAD/FAD binding subunit | -3.04 |
| **Propanoate metabolism** | | | |
| 0990 | mgsA | methylglyoxal synthase | 6.02 |
| 0687 | plfB | formate C-acetyltransferase | -2.24 |
| **Carbon Fermentation** | | | |
| 2070 | ldh | D-lactate dehydrogenase | 2.68 |
| 1007 | | nickel-dependent lactate racemase | 2.43 |
| 1359 | | lactate utilization protein | 2.84 |
| 2233 | | putative transport protein (AspT/YidE/YbjL antiporter duplication domain protein) | -2.29 |
| 2880 | | putative acyl-CoA reductase/dehydratase | -2.80 |
| **Sugar transport systems** | | | |
| 0304 | rbsA | ribose transport system ATP-binding protein | 4.45 |
| 0303 | rbsB | D-ribose ABC transporter, substrate-binding protein | 3.90 |
| 0305 | rbsC | ribose ABC transporter, permease protein | 6.84 |
| 2927 | | cellobiose-phosphate degrading protein | 2.40 |
| 2928 | | PTS system, N-acetylglucosamine-specific IIB component | 2.38 |
| 0210 | | putative sugar-phosphate kinase | 2.84 |
| 2895 | | PTS system, alpha-glucoside-specific IIBC component | -39.14 |
| 2554 | crr | PTS system, glucose-specific IIa component | -3.02 |
| 2223 | mtlA | PTS system, mannitol-specific IIBC component | -2.90 |
| 2436 | | multiple sugar transport system permease protein | -3.10 |
| **Unknown pathway** | | | |
| 1055 | | glycosyltransferase family 2 protein | 12.57 |

| | | | |
|---|---|---|---|
| 2393 | | Cof-type HAD-IIB family hydrolase | 4.66 |
| 1405 | | putative polysaccharide deacetylase | 2.92 |
| 2958 | | putative glycosyltransferase | -2.86 |

### Nitrogen Metabolism

#### Glycine, serine and threonine metabolism

| | | | |
|---|---|---|---|
| 0656 | gcvH | glycine cleavage system H protein | 2.58 |
| 1556 | gcvPB | glycine dehydrogenase subunit 2 | 2.47 |
| 2615 | glyA_1 | glycine hydroxymethyltransferase | 2.10 |
| 2406 | tdcB | threonine dehydratase | 2.63 |
| 2026 | thrB | homoserine kinase | 3.21 |

#### Cysteine metabolism

| | | | |
|---|---|---|---|
| 3434 | | 5-methyltetrahydrofolate--homocysteine methyltransferase | 3.01 |
| 1384 | sseA | thiosulfate/3-mercaptopyruvate sulfurtransferase | 2.24 |
| 3263 | | cysteine-S-conjugate beta-lyase | 21.92 |
| 3436 | luxS | S-ribosylhomocysteine lyase | 3.49 |
| 2078 | | S-methylcysteine transport system ATP-binding protein | -2.07 |
| 2377 | | cysteine-S-conjugate beta-lyase | -6.96 |

#### Alanine, aspartate and glutamate metabolism

| | | | |
|---|---|---|---|
| 0307 | | Cytoplasmic protein (GATase1_like) | 4.97 |
| 1386 | aspB | glutamate synthase (NADPH) small chain | 2.05 |
| 2719 | | aspartate aminotransferase | 2.18 |

#### Arginine biosynthesis

| | | | |
|---|---|---|---|
| 0306 | argE | acetylornithine deacetylase | 2.66 |
| 0014 | | Protein Arginine kinase | -2.66 |

#### Lysine Biosynthesis

| | | | |
|---|---|---|---|
| 3081 | | N-acetyldiaminopimelate deacetylase | 2.88 |
| 3086 | dapB1 | 4-hydroxy-tetrahydrodipicolinate reductase | -2.40 |

#### Stickland reactions (amino acid fermentation)

| | | | |
|---|---|---|---|
| 3097 | prdF | proline racemase | 2.06 |
| 3099 | prdE | D-proline reductase (dithiol)-stabilizing protein | 2.60 |
| 2240 | grdA | glycine/sarcosine/betaine reductase complex component A | -2.22 |
| 1588 | trxA1 | thioredoxin 1 | -2.08 |
| 1589 | trxB1 | thioredoxin reductase (NADPH) | -2.55 |

#### Peptide degradation/amino acid production

| | | | |
|---|---|---|---|
| 0931 | | putative aminopeptidase | 2.12 |
| 3199 | | putative nitroreductase | 3.17 |
| 1242 | | D-stereospecific aminopeptidase | 2.32 |
| 2704 | | L-2-amino-thiazoline-4-carboxylic acid hydrolase | 2.49 |
| 2878 | | transglutaminase domain-containing protein (ChW repeat-/cell adhesion) | 2.70 |
| 0308 | abgB1 | aminobenzoyl-glutamate utilization protein B | 4.67 |
| 3326 | | oligoendopeptidase F | 5.55 |
| 3433 | | aminopeptidase | 4.19 |
| 0089 | map1 | methionyl aminopeptidase | -2.78 |
| 0957 | | nitroreductase-family protein | -2.47 |
| 2072 | msrAB | peptide methionine sulfoxide reductase msrA/msrB | -2.97 |
| 1991 | | putative amidohydrolase (YgeY family selenium metabolism-linked hydrolase) | -2.99 |

#### Nitrogen source uptake

| | | | |
|---|---|---|---|
| 2590 | brnQ | putative branched-chain amino acid transport system II carrier protein | 3.28 |
| 1099 | | branched-chain amino acid transport system II carrier protein (LIVCS family) | 2.82 |
| 1594 | | alanine or glycine:cation symporter, AGCS family | -2.27 |
| 2161 | | proton-dependent oligopeptide transporter, POT family | -2.66 |
| 2871 | | proton-dependent oligopeptide transporter | -3.88 |

#### Unknown pathway

| | | | |
|---|---|---|---|
| 1857 | | amino acid-binding protein (ACT domain-containing protein) | 5.56 |
| 2733 | | nitrilase | 2.47 |
| 2074 | hcp | hydroxylamine reductase | -3.05 |
| 1521 | | putative nitric oxide reductase flavoprotein (NADH oxidase (H2O-forming) | -2.19 |

### Nucleic acid metabolism

#### Purine metabolism

| | | | |
|---|---|---|---|
| 1058 | nudF | ADP-ribose pyrophosphatase | 3.42 |
| 1061 | deoB | phosphopentomutase | 4.04 |
| 2224 | guaB | IMP (inosine-5'-monophosphate) dehydrogenase | 2.06 |
| 0618 | | phosphoribosyl 1,2-cyclic phosphate phosphodiesterase | -3.91 |
| 0088 | adk | adenylate kinase | -2.61 |

#### Pyrimidine metabolism

| | | | |
|---|---|---|---|
| 3313 | | dCMP deaminase | 2.32 |
| 0185 | pyrB | aspartate carbamoyltransferase catalytic subunit | 2.20 |
| 3429 | pyrAA2 | carbamoyl-phosphate synthase small subunit | -2.03 |
| 2045 | pyrH | uridylate kinase | -2.17 |
| 1819 | | dCMP deaminase | -3.77 |
| 0196 | | pyridinium-3,5-biscarboxylic acid mononucleotide synthase | -3.35 |
| 0711 | | cytidine/deoxycytidylate deaminase family protein | -7.51 |

#### Nucleotide degradation

| | | | |
|---|---|---|---|
| 1129 | | ribonuclease J | 2.05 |
| 1238 | | RidA family protein (putative endoribonuclease) | 2.21 |
| 0490 | nth | endonuclease III | -2.28 |
| 3169 | rph | ribonuclease Ph | -2.34 |
| 3232 | uvrC | excinuclease ABC subunit C | -3.04 |
| 3522 | ssb | single-strand DNA-binding protein | -2.29 |
| 0485 | nfo | deoxyribonuclease IV | -4.16 |
| 0042 | | ribonuclease III family protein | -2.11 |
| 0013 | | putative DNA repair protein/DNA methylase/uvrB/C motif family protein | -2.41 |

#### Unknown pathway

| | | | |
|---|---|---|---|
| 0615 | | putative nucleotide phosphodiesterase | 2.12 |
| 1143 | | nucleotide pyrophosphohydrolase | 3.38 |
| 1410 | | xanthine dehydrogenase accessory factor | 2.82 |
| 3235 | | putative nucleotide pyrophosphatase | -3.86 |
| 0618 | | phosphoribosyl 1,2-cyclic phosphate phosphodiesterase | -3.91 |

### Energy generation

#### ATP synthases

| | | | |
|---|---|---|---|
| 3309 | atpE | F-type H+-transporting ATPase subunit c | 2.21 |
| 2789 | ntpA | V/A-type H+/Na+-transporting ATPase subunit A | 2.43 |
| 2788 | ntpB | V/A-type H+/Na+-transporting ATPase subunit B | 3.30 |
| 2791 | ntpC | V/A-type H+/Na+-transporting ATPase subunit C | 2.87 |
| 2787 | ntpD | V/A-type H+/Na+-transporting ATPase subunit D | 2.52 |
| 2792 | ntpE | V/A-type H+/Na+-transporting ATPase subunit E | 3.33 |
| 2794 | ntpI | V/A-type H+/Na+-transporting ATPase subunit I | 2.47 |
| 2793 | ntpK | V/A-type H+/Na+-transporting ATPase subunit K | 2.25 |
| 2795 | ntpG | V/A-type H+/Na+-transporting ATPase subunit G/H | 2.59 |

#### Electron transport

| | | | |
|---|---|---|---|
| 0912 | etfA2 | electron transfer flavoprotein alpha subunit | 2.04 |
| 1009 | etfA3 | electron transfer flavoprotein alpha subunit | 3.33 |
| 0911 | etfB2 | electron transfer flavoprotein beta subunit | 2.71 |
| 1008 | etfB3 | electron transfer flavoprotein beta subunit | 2.91 |
| 2907 | | energy-coupling factor transport system substrate-specific component | 3.64 |
| 0735 | | electron transfer flavoprotein beta subunit | -4.03 |

### Translation

#### Ribosomal proteins

| | | | |
|---|---|---|---|
| 2360 | lepA | GTP-binding protein LepA (elongation factor EF-4) | 2.28 |
| 2366 | rpsT | small subunit ribosomal protein S20 | -2.18 |
| 0070 | rplB | large subunit ribosomal protein L2 | -3.28 |
| 0067 | rplC | large subunit ribosomal protein L3 | -2.21 |
| 0068 | rplD | large subunit ribosomal protein L4 | -2.68 |
| 0082 | rplF | large subunit ribosomal protein L6 | -2.45 |
| 0102 | rplM | large subunit ribosomal protein L13 | -2.04 |
| 0086 | rplO | large subunit ribosomal protein L15 | -2.24 |
| 0074 | rplP | large subunit ribosomal protein L16 | -2.56 |
| 0083 | rplR | large subunit ribosomal protein L18 | -4.99 |
| 0069 | rplW | large subunit ribosomal protein L23 | -2.71 |
| 0075 | rpmC | large subunit ribosomal protein L29 | -6.39 |
| 0085 | rpmD | large subunit ribosomal protein L30 | -3.42 |
| 0612 | rpmI | large subunit ribosomal protein L35 | -2.23 |
| 2047 | rpsB | small subunit ribosomal protein S2 | -2.38 |
| 0073 | rpsC | small subunit ribosomal protein S3 | -2.43 |
| 0084 | rpsE | small subunit ribosomal protein S5 | -3.35 |
| 0091 | infA | translation initiation factor IF-1 | -4.53 |
| 2046 | tsf | translation initiation factor IF-1 | -2.02 |

#### Ribosome biogenesis

| | | | |
|---|---|---|---|
| 0916 | | 23S rRNA pseudouridine1911/1915/1917 synthase | 2.74 |

| | | | |
|---|---|---|---|
| 1768 | | serine/arginine repetitive matrix protein 2 | 3.86 |
| 2329 | era | GTPase | 2.19 |
| 3161 | engB | ribosome biogenesis GTP-binding protein | 3.59 |
| 2550 | | ribosome-binding ATPase | 10.09 |
| 1095 | rimM | 16S rRNA processing protein | -5.53 |
| 1138 | rimP | ribosome maturation factor | -3.17 |
| 3508 | | 23S rRNA (pseudouridine1915-N3)-methyltransferase | -4.77 |
| **tRNA synthesis and processing** | | | |
| 1966 | glnS | glutaminyl-tRNA synthetase | 2.76 |
| 2144 | asnC | asparagine--tRNA ligase | 2.36 |
| 1892 | | Ala-tRNA(Pro) deacylase | 2.47 |
| 0038 | proS | prolyl-tRNA synthetase | -2.40 |
| 3536 | trmE | tRNA modification GTPase | -2.28 |
| 0491 | | tRNA (cytidine/uridine-2'-O-)-methyltransferase | -3.47 |
| 2372 | | tRNA threonylcarbamoyladenosine dehydratase | -4.39 |
| **Chaperone** | | | |
| 2148 | | Hypothetical protein (VWA domain-containing protein) | 2.27 |
| 2149 | | MoxR family ATPase | 2.89 |
| 0336 | ppiB | peptidyl-prolyl cis-trans isomerase B | 3.10 |
| 2353 | dnaJ | molecular chaperone | -2.26 |
| 2354 | dnaK | molecular chaperone | -2.49 |
| 0195 | groEL | 60 kDa chaperonin | -2.58 |
| 2355 | grpE | molecular chaperone | -2.44 |
| 1933 | clpB | ATP-dependent Clp protease ATP-binding subunit ClpB | -1.93 |
| 0015 | clpC | ATP-dependent Clp protease ATP-binding subunit ClpC | -2.33 |
| **Toxin production** | | | |
| 0584 | tcdA | toxin A/B | 1.87 |
| 0582 | tcdB | toxin A/B | 2.42 |
| **Sporulation and germination** | | | |
| **Sporulation pathway** | | | |
| 3532 | soj | sporulation initiation inhibitor | 2.70 |
| 0700 | spoIIAB | stage II sporulation protein AB (anti-sigma F factor) | 3.88 |
| 1052 | spo0A | stage 0 sporulation protein A | 1.81 |
| 0699 | spoIIAA | stage II sporulation protein AA | 2.71 |
| 0714 | spoIVB | stage IV sporulation protein | -4.00 |
| 1001 | obg | Spo0B-associated GTP-binding protein | -2.92 |
| 3335 | spoIIIF | stage V sporulation protein B | -2.53 |
| 3353 | spoVG | stage V sporulation protein G | -2.21 |
| **Germination** | | | |
| 0476 | sleC | spore cortex-lytic enzyme pre-pro-form | -9.72 |
| **Membrane biogenesis and cell surface proteins** | | | |
| **Fatty acid biogenesis** | | | |
| 2497 | | DegV (Bs) lipid transport or fatty acid metabolism | 2.99 |
| 0918 | acpP | acyl carrier protein | 3.07 |
| 1859 | accA | acetyl-CoA carboxylase carboxyl transferase subunit alpha | 3.03 |
| 1017 | fabH | 3-oxoacyl-[acyl-carrier-protein] synthase III | -2.07 |
| 0059 | fabG | NADP-dependent 7-alpha-hydroxysteroid dehydrogenase | -2.80 |
| 1836 | eutL | ethanolamine utilization protein | -10.25 |
| **Peptidoglycan metabolism** | | | |
| 2656 | | putative cell-wall hydrolase (NlpC/P60 family protein) | 2.63 |
| 2390 | | serine-type D-Ala-D-Ala carboxypeptidase | 3.26 |
| 1255 | ddl | D-alanine-D-alanine ligase | -3.17 |
| 1463 | | N-acetylmuramoyl-L-alanine amidase | -5.56 |
| 0866 | nagA | N-acetylglucosamine-6-phosphate deacetylase | -2.34 |
| 0867 | glmD | glucosamine-6-phosphate deaminase | -3.70 |
| 2662 | | putative teichuronic acid biosynthesis glycosyl transferase | -2.18 |
| **Cell surface proteins** | | | |
| 0891 | cwp16 | N-acetylmuramoyl-L-alanine amidase | 2.56 |
| 2679 | | PIG-L family deacetylase | 2.58 |
| 2676 | cwp84 | cell surface-associated cysteine protease | 2.33 |
| 1318 | cwp20 | putative penicillin-binding protein | 2.57 |
| 1911 | cwp28 | cell wall-binding protein | 5.32 |
| 2099 | | cell surface protein | 3.38 |
| 2194 | | putative membrane protein precursor | -2.27 |

##### Protein export

| | | | |
|---|---|---|---|
| 2508 | hlyD | secretion protein HlyD | 2.76 |
| 0087 | prlA | preprotein translocase subunit SecY | -3.88 |

#### Other transport systems

##### Iron uptake systems

| | | | |
|---|---|---|---|
| 0973 | rnfC | Na+-translocating ferredoxin:NAD+ oxidoreductase subunit C | 2.28 |
| 3447 | | Fe-S cluster assembly ATP-binding cassette domain-containing protein | 2.95 |
| 2271 | fhuC | iron complex transport system permease protein | -3.89 |
| 2273 | fhuB | iron complex transport system ATP-binding protein | -3.16 |
| 2774 | fhuD | iron complex transport system substrate-binding protein | -2.51 |
| 1326 | | ferrous iron transport protein A | -11.82 |
| 1639 | | ferrous iron transport protein A | -2.26 |
| 1328 | feoB1 | ferrous iron transport protein B | -2.21 |
| 3134 | feoA3 | ferrous iron transport protein A | -2.79 |

##### ABC transporters

| | | | |
|---|---|---|---|
| 1376 | | putative ABC transport system permease protein | 2.65 |
| 1377 | | putative ABC transport system ATP-binding protein | 3.21 |
| 0873 | | ATP-binding cassette | -6.24 |
| 0874 | | putative ABC transporter, permease/ATP-binding protein | -6.01 |
| 0716 | | ABC transporter, ATP-binding protein | -2.22 |
| 0799 | modB | molybdate transport system permease protein | -2.38 |
| 2010 | | putative ABC transporter, permease protein | -3.84 |
| 2117 | | ATP-binding cassette, subfamily B, bacterial | -4.37 |
| 2118 | | ATP-binding cassette, subfamily B, bacterial | -4.42 |
| 2825 | | sulfonate transport system ATP-binding protein | -4.10 |
| 2826 | | sulfonate transport system permease protein | -2.90 |

##### Symporters

| | | | |
|---|---|---|---|
| 2109 | | sodium:solute symporter family protein | 2.80 |
| 2515 | | glutamate:GABA antiporter | 3.40 |
| 2233 | | AspT/YidE/YbjL antiporter duplication domain protein | -2.29 |
| 2077 | | putative Sodium:dicarboxylate symporter | -4.16 |
| 2138 | | solute:Na+ symporter, SSS family | -2.58 |
| 2557 | | putative Na(+)/H(+) antiporter | -8.62 |
| 2770 | | MATE family efflux transporter | -3.48 |

##### Other systems

| | | | |
|---|---|---|---|
| 1308 | | pyridoxamine 5'-phosphate oxidase family protein (MFS transporter) | 2.36 |
| 2931 | | putative amino acid permease | 3.10 |
| 3096 | | permease (sulfite exporter TauE/SafE family protein) | 2.42 |
| 1333 | ssuA | sulfonate transport system substrate-binding protein | 2.23 |
| 2743 | dltB | membrane protein involved in D-alanine export | 2.71 |
| 1779 | | multidrug transporter MatE | -4.64 |

#### Cell surface organelles

##### Type IV pillus biogenesis

| | | | |
|---|---|---|---|
| 3342 | pilT | twitching motility protein PilT | 2.39 |
| 3345 | pilV | Type IV pilin | 5.13 |
| 3346 | pilO | putative type IV pilus assembly protein | 2.60 |
| 3350 | pilA1 | Pilin subunit | 3.18 |

##### Flagella biogenesis

| | | | |
|---|---|---|---|
| 0264 | fliP | flagellar biosynthetic protein | -2.40 |
| 0275 | fliN | flagellar motor switch protein | -3.29 |
| 0253 | fliH | flagellar assembly protein | -3.19 |
| 0265 | fliQ | flagellar biosynthetic protein | -1.81 |
| 0270 | fliA | RNA polymerase sigma factor for flagellar operon | -1.63 |

#### Regulation

##### Two component systems

| | | | |
|---|---|---|---|
| 1194 | | two-component system, OmpR family | 2.16 |
| 1635 | | two-component sensor histidine kinase | 3.14 |
| 2760 | | two-component system, sensor histidine kinase | 6.02 |
| 1748 | | two-component response regulator | -36.61 |
| 1749 | | two-component response regulator | -4.60 |
| 0534 | | two-component response regulator | -2.81 |
| 0869 | | two-component response regulator | -2.82 |
| 1872 | | putative two-component sensor histidine kinase | -2.22 |
| 3126 | | two-component response regulator | -4.90 |

#### HTH regulators

| | | | |
|---|---|---|---|
| 1278 | | bacterial regulatory helix-turn-helix, AraC family protein | 3.94 |
| 2872 | | CarD family transcriptional regulator | 2.98 |
| 0552 | | CRP/FNR family transcriptional regulator, anaerobic regulatory protein | 6.19 |
| 2184 | | GntR-family transcriptional regulator (PLP-dependent aminotransferase family protein) | 3.11 |
| 0817 | | GntR-family transcriptional regulator | 3.19 |
| 1105 | | CarD family transcriptional regulator | 2.07 |
| 1572 | | helix-turn-helix transcriptional regulator | 2.96 |
| 2110 | | TetR-family transcriptional regulator | 3.75 |
| 3046 | | MerR-family transcriptional regulator | 11.46 |
| 3379 | | Lrp/AsnC family transcriptional regulator, leucine-responsive regulatory protein | 2.30 |
| 2929 | treR | GntR family transcriptional regulator, trehalose operon transcriptional repressor | 3.51 |
| 0301 | rbsR | LacI family transcriptional regulator | 2.21 |
| 1006 | | transcriptional repressor for pyruvate dehydrogenase complex | -2.74 |
| 1590 | | ArsR-family transcriptional regulator | -3.01 |
| 2356 | hrcA | heat-inducible transcriptional repressor | -2.72 |
| 2653 | | LytR family transcriptional regulator | -3.43 |

#### Transcription

| | | | |
|---|---|---|---|
| 0096 | rpoA | DNA-directed RNA polymerase subunit alpha | -2.53 |

#### Antiterminator

| | | | |
|---|---|---|---|
| 3324 | rho | transcription termination factor Rho | 2.10 |
| 0206 | | putative transcription antiterminator | 3.36 |
| 2556 | licT | beta-glucoside operon transcriptional antiterminator | -2.32 |

#### Stress response

| | | | |
|---|---|---|---|
| 0822 | cspA | cold shock protein | 5.75 |
| 1197 | cspB | cold shock protein | 2.26 |
| 2345 | | CRISPR-associated protein Cas5 | 2.36 |
| 2447 | | Asp23/Gls24 family envelope stress response protein | 3.53 |
| 0012 | ctsR | transcriptional regulator of stress and heat shock response | -2.88 |

#### Signalling proteins

| | | | |
|---|---|---|---|
| 2076 | | cyclic nucleotide (cAMP?)-binding domain protein | 2.77 |
| 1267 | dccA | diguanylate cyclase | 28.15 |
| 2799 | | putative phosphodiesterase | 2.77 |
| 1591 | | putative diguanylate cyclase | -2.12 |

#### Putative biofilm regulators

| | | | |
|---|---|---|---|
| 2121 | sinR | putative regulatory protein | -3.29 |
| 2122 | | putative regulatory protein | 2.54 |

### Cofactor metabolism

#### Ferredoxins, iron-sulphur proteins

| | | | |
|---|---|---|---|
| 2059 | | iron-only hydrogenase system regulator | 2.31 |
| 2101 | | ferritin | 3.15 |
| 2126 | | putative radical SAM superfamily lipoprotein | 2.60 |
| 1667 | | Hypothetical protein (SufBD protein) | -2.77 |
| 2736 | rbr | rubrerythrin | -3.99 |
| 1925 | fldX | flavodoxin | -4.53 |

#### Other cofactors

| | | | |
|---|---|---|---|
| 2413 | nadD | nicotinate-nucleotide adenylyltransferase | 2.57 |
| 3261 | cobT | nicotinate-nucleotide--dimethylbenzimidazole phosphoribosyltransferase | 2.20 |
| 3246 | cbiH | precorrin-3B C17-methyltransferase | 4.52 |
| 3244 | cbiK | sirohydrochlorin cobaltochelatase | 4.33 |
| 1510 | | GTP 3',8-cyclase | 2.38 |
| 1876 | | F420-0--gamma-glutamyl ligase | 2.85 |
| 1383 | | gamma-glutamyltranspeptidase / glutathione hydrolase | 2.28 |

### Cell division

| | | | |
|---|---|---|---|
| 0627 | zapA | cell division protein | 2.29 |
| 1983 | | chromosome segregation protein | 2.02 |
| 2506 | sepF | cell division inhibitor | 2.01 |
| 0988 | minE | cell division topological specificity factor | 2.23 |

### Toxin/antitoxin system

| | | | |
|---|---|---|---|
| 1759 | | addiction module toxin, rele/stbe family | -6.85 |
| 1760 | | addiction module antitoxin, relb/dinj family | -7.17 |

### Mobile elements

#### Prophage island 1

| | | | |
|---|---|---|---|
| 1419 | | phage antirepressor protein | -2.80 |
| 1428 | | prophage antirepressor (Bro-N domain-containing protein) | -5.00 |
| 1429 | | hypothetical protein | -2.50 |
| 1431 | | putative phage DNA-binding protein | -5.54 |
| 1432 | | hypothetical phage protein | -4.02 |
| 1433 | | hypothetical phage protein | -3.41 |
| 1434 | | hypothetical phage protein | -4.30 |
| 1435 | | phage protein | -4.81 |
| 1436 | | phage protein | -3.75 |
| 1437 | | hypothetical phage protein | -2.81 |
| 1438 | | phage protein | -2.99 |
| 1439 | | phage protein | -2.72 |
| 1440 | | phage protein | -4.10 |
| 1441 | | phage protein | -5.09 |
| 1442 | | phage protein | -5.47 |
| 1443 | | hypothetical phage protein | -3.30 |
| 1444 | | hypothetical phage protein | -5.86 |
| 1445 | | phage protein | -4.80 |
| 1446 | | phage protein | -3.62 |
| 1448 | | hypothetical phage protein | -3.81 |
| 1449 | | hypothetical phage protein | -3.92 |
| 1450 | | hypothetical phage protein | -3.23 |
| 1451 | | hypothetical phage protein | -4.11 |
| 1452 | | phage protein | -2.91 |
| 1454 | | putative uncharacterized protein | -5.53 |
| 1455 | | phage protein | -4.38 |
| 1456 | | phage protein | -6.10 |
| 1457 | | hypothetical phage protein | -4.93 |
| 1458 | | putative uncharacterized protein | -4.42 |
| 1459 | | hypothetical phage protein | -5.01 |
| 1460 | | phage protein | -5.01 |
| 1462 | | putative uncharacterized protein | -2.00 |

##### Putative phage island

| | | | |
|---|---|---|---|
| 1205 | | hypothetical protein | -3.49 |
| 1206 | | phage tail sheath protein | -3.97 |
| 1207 | | Phage portal protein | -3.09 |
| 1208 | | Phage portal protein | -3.68 |
| 1209 | | hypothetical protein | -3.48 |
| 1210 | | putative phage protein | -2.94 |
| 1211 | | putative phage protein | -2.96 |
| 1212 | | putative phage cell wall hydrolase | -3.03 |
| 1213 | | hypothetical protein | -3.66 |
| 1214 | | putative phage protein | -2.81 |
| 1215 | | putative phage protein | -3.27 |
| 1216 | | putative phage protein | -3.11 |
| 1217 | | putative phage tail fiber protein | -3.29 |
| 1218 | | putative phage protein | -2.77 |
| 1219 | | putative phage-related protein | -4.88 |
| 1220 | | putative membrane protein | -2.26 |

##### Conjugation proteins

| | | | |
|---|---|---|---|
| 1797 | | ATP-dependent endonuclease (putative conjugative transposon DNA recombination protein) | 2.33 |
| 1798 | | helicase UvrD (putative conjugative transposon conserved hypothetical protein) | 2.20 |
| 1799 | | conjugal transfer protein (transposase) | 2.79 |
| 1747 | | putative conjugative transposon regulatory protein | -8.75 |
| 2959 | tndX | conjugative transposon site-specific recombinase | -4.18 |

##### Plasmid replication

| | | | |
|---|---|---|---|
| 1935 | | plasmid replication protein | -6.59 |

#### Function unknown

##### Putative membrane, lipoproteins or exported proteins

| | | | |
|---|---|---|---|
| 0706 | | putative membrane protein | 2.68 |
| 0707 | | putative membrane protein | 3.77 |
| 0471 | | putative membrane protein | 2.19 |
| 0760 | | putative membrane protein | 3.13 |
| 1283 | | putative membrane protein | 3.02 |
| 1290 | | YitT family protein (putative membrane protein) | 2.65 |
| 1346 | | putative exported protein | 3.00 |
| 1663 | | putative membrane protein | 2.55 |

| | | |
|---|---|---|
| 1815 | putative membrane protein (YtxH domain-containing protein) | 7.05 |
| 2373 | putative exported protein | 2.58 |
| 2536 | putative exported protein | 3.38 |
| 0947 | putative membrane protein | -7.24 |
| 1464 | cell surface protein | -2.40 |
| 2055 | putative exported protein (VanW-llike protein) | -2.60 |
| 2686 | adhesion protein | -3.46 |
| 3168 | Metallophosphoesterase | -2.64 |
| Other | | |
| 0904 | putative phosphoesterase | 3.66 |
| 0532 | GrpB family protein | 8.69 |
| 1325 | putative hydrolase | 3.91 |
| 3431 | conserved hypothetical protein (putative zinc-binding protein) | 3.56 |
| 0708 | putative amidohydrolase | 2.66 |
| 1347 | hypothetical protein | 2.96 |
| 0147 | conserved hypothetical protein | 3.17 |
| 0504 | conserved hypothetical protein (hydrolase) | 3.05 |
| 1657 | conserved hypothetical protein | 8.49 |
| 1536 | conserved hypothetical protein | 2.24 |
| 2039 | conserved hypothetical protein | 2.73 |
| 3525 | conserved hypothetical protein | 6.79 |
| 2005 | hypothetical protein | 2.65 |
| 2123 | hypothetical protein | 5.89 |
| 2499 | hypothetical protein | 4.46 |
| 0926 | hypothetical protein | 8.76 |
| 0961 | uncharacterized protein | 10.40 |
| 3010 | hypothetical protein | 2.36 |
| 0489 | putative ATP-binding protein | -2.01 |
| 0049 | NYN domain-containing protein | -3.13 |
| 1672 | Hypothetical protein (arsenate reductase family protein) | -3.85 |
| 1772 | putative uncharacterized protein (recombinase) | -4.75 |
| 1780 | putative uncharacterized protein (helix-turn-helix domain-containing protein) | -5.00 |
| 2369 | conserved hypothetical protein (GNAT family N-acetyltransferase) | -2.21 |
| 0828 | conserved hypothetical protein | -2.04 |
| 0875 | conserved hypothetical protein | -18.87 |
| 2370 | conserved hypothetical protein | -2.53 |
| 2873 | conserved hypothetical protein | -2.16 |
| 2874 | conserved hypothetical protein (ATP-binding protein) | -3.82 |
| 2875 | conserved hypothetical protein | -3.59 |
| 3452 | conserved hypothetical protein | -6.37 |
| 1025 | conserved hypothetical protein | -16.61 |
| 1138 | conserved hypothetical protein | -7.27 |
| 1717 | conserved hypothetical protein | -3.74 |
| 1788 | putative uncharacterized protein | -11.36 |
| 2988 | putative uncharacterized protein | -8.17 |
| 1771 | putative uncharacterized protein | -2.28 |
| 1773 | putative uncharacterized protein | -5.88 |
| 1777 | putative uncharacterized protein | -6.66 |
| 1934 | hypothetical protein | -5.78 |
| 1954 | hypothetical protein | -3.61 |
| 2250 | hypothetical protein | -2.41 |
| 2300 | hypothetical protein | -15.63 |
| 2391 | hypothetical protein | -5.31 |
| 3448 | hypothetical protein | -2.60 |

[1]Gene number CDR20291_
