## Supplemental Table 4 for "Insights into the regulatory mechanisms of *Clostridioides difficile* biofilm formation"

| Start | Finish | Genes flanking | Length (bp) | Comparison with 630 genome |
|---|---|---|---|---|
| 17395 | 17529 | CDR20291_0002 & SerS1 | 155 | CDs001 T-box leader (17634-17855). Early termination of transcript due to stop codon compared with strain 630 (222 bp). |
| 19880 | 20166 | CDR20291_t11 & dnaH | 287 | CDs002 SRP bact RNA (20243-20343). |
| 72841 | 72982 | CDR20291_ispF & proS | 162 | CDs003 T-box leader (73019-73242). Early termination of transcript due to stop codon compared with strain 630 (224 bp). |
| 169680 | 169798 | CDR20291_0128 & metE | 111 | CDs005 S box leader (174174-174250). |
| 175839 | 176092 | CDR20291_0132 & 0133 | 254 | CDR20291_CDs001. Loci present in strain 630 (180306-180559). |
| 192241 | 192428 | CDR20291_0145 & 0146 | 188 | CDs006 RFN element (196732-196849). |
| 251813 | 251915 | CDR20291_0197 & guaA | 103 | CDs007 Purine riboswitch (256217-256319). |
| 308009 | 308223 | CDR20291_0247 & flgB | 212 | Cdi1_3 c-di-GMP riboswitch (308776-309272). |
| 368784 | 368912 | CDR20291_0310 & 0311 | 129 | CDR20291_CDs002. Loci not present in strain 630. |
| 392409 | 392601 | CDR20291_0328 & cbiM | 193 | CDs008 Cobalamin riboswitch (391011-391203). |
| 400935 | 401068 | ppiB & CDR20291_0337 | 134 | CDR20291_CDs003. Loci present in strain 630. |
| 488181 | 488363 | CDR20291_0404 & 0405 | 183 | CDR20291_CDs004. Loci not present in strain 630. |
| 489730 | 489934 | CDR20291_0405 & 0406 | 205 | CDR20291_CDs005. Loci not present in strain 630. |
| 599578 | 599833 | thrS & CDR20291_0497 | 256 | CDs016 T-box leader (683173-683427). |
| 620313 | 620597 | CDR20291_0512 & 0513 | 285 | CDR20291_CDs006. Loci partially present in strain 630 (94% identity) (705355-705571). |
| 684561 | 684762 | CDR20291_0564 & 0565 | 202 | CDR20291_CDs007. Loci not present in strain 630. |
| 755233 | 755393 | CDR20291_0610 & infC | 161 | CDR20291_CDs008. Loci present in strain 630. |
| 772939 | 773189 | CDR20291_0624 & pheS | 251 | CDs018 T-box leader (849440-849690). |
| 841259 | 841421 | CDR20291_0677 & 0678 | 160 | CDs019 T-box leader (917371-917598). |
| 865921 | 866081 | CDR20291_0697 & 0698 | 161 | CDR20291_CDs009. Loci not present in strain 630. |
| 978000 | 978189 | CDR20291_0804 & 0805 | 223 | CD630_n00330/sCD1053.1 ncRNA IGR (1052640-1052823). |
| 981007 | 981380 | CDR20291_0807 & 0808 | 304 | CDR20291_CDs010. Loci present in strain 630. |
| 983518 | 983797 | CDR20291_0808 & tlpB | 280 | CDR20291_CDs011. Loci present in strain 630. |
| 985245 | 985370 | tlpB & CDR20291_0810 | 126 | CDR20291_CDs012. Loci not present in strain 630. |
| 1025919 | 1026192 | CDR20291_0841 & leuA | 263 | CDs021 T-box leader (1152624-1152886). |
| 1138657 | 1138849 | CDR20291_0925 & 0926 | 193 | CD630_n00410 (RCd3 6S RNA) (1259900-1260092). |
| 1173955 | 1174314 | CDR20291_0958 & 0959 | 360 | CDR20291_CDs013. Loci not present in strain 630. Rfam homology to Group I introns |
| 1175712 | 1175834 | CDR20291_0959 & 0960 | 123 | CDR20291_CDs014. Loci not present in strain 630. CDR_0959 is tlpB |
| 1219842 | 1219948 | CDR20291_1002 & hom1 | 107 | CDs022 S-box leader (1364065-1364171). |
| 1289619 | 1289869 | CDR20291_1070 & 1071 | 251 | CDR20291_CDs015. Loci not present in strain 630. |
| 1291581 | 1293267 | CDR20291_1071 & 1072 | 1687 | CD630_n00460 CRIPSR 6 (1434587-1435639). |
| 1346436 | 1346689 | trmU & alaS | 254 | CDs023 T-box leader (Ala) (1488489-1488742). |
| 1501290 | 1502390 | CDR20291_1265 & 1266 | 1101 | CD630_n00510 CRISPR 7 (1645024-1646018). |
| 1580675 | 1581025 | CDR20291_1337 & 1338 | 351 | CDR20291_CDs016. Loci not present in strain 630. Rfam homology to Group I introns |
| 1582788 | 1582894 | CDR20291_1338 & metN | 107 | CDs025 S box leader (1724139-1724245). |
| 1612906 | 1613106 | CDR20291_1364 & 1365 | 201 | CD630_n00560 CRISPR 8 (1756109-1756833). |
| 1617414 | 1617650 | feoA2 & CDR20291_1368 | 237 | SQ1002 ncRNA IGR (1760892-1761128). |
| 1619411 | 1619679 | CDR20291_1369 & tyrR | 269 | CDs027/SQ1005 T-box leader (1762889-1763157). |
| 1667086 | 1667374 | CDR20291_1411 & 1412 | 289 | CDs028 T-box leader (1810573-1810861). |
| 1705582 | 1705684 | CDR20291_1445 & 1446 | 100 | CDR20291_CDs017. Loci not present in strain 630. |
| 1721530 | 1721661 | CDR20291_1461 & 1462 | 132 | CDR20291_CDs018. Loci not present in strain 630. Rfam homology abiF RNA motif |
| 1743469 | 1743955 | CDR20291_1476 & hom2 | 257 | CDs030 T-box leader (1830948-1831204). |
| 1743775 | 1744026 | CDR20291_1476 & hom2 | 252 | CDs031 T-box leader (1831254-1831505). |
| 1767068 | 1767161 | CDR20291_1496 & thiD | 94 | CDs032 TPP riboswitch (THI element) (854584-1854677). |
| 1829965 | 1830070 | CDR20291_1551 & IplA | 106 | CDs033 S-box leader (1917499-1917604). |
| 1831312 | 1831488 | IplA & CDR20291_1553 | 177 | CDs034 lysine riboswitch (1918846-1919022). |
| 1834460 | 1834654 | CDR20291_1554 & 1555 | 195 | CDR20291_CDs019. Loci present in strain 630. |
| 1845366 | 1845528 | guaD & CDR20291_1561 | 162 | CD630_n00600 CRISPR 9 (1935320-1935481). |
| 1853237 | 1853454 | CDR20291_1564 & 1565 | 218 | CD630_n00620 ncRNA IGR (1943181-1943450). |
| 1853584 | 1853755 | CDR20291_1564 & 1565 | 172 | CDR20291_CDs020. Loci present in strain 630 (1943529-1943700). |
| 1883259 | 1883380 | ribD & recQ | 123 | CDs036 RFN/ FMN element (1973231-1973352c). |
| 1886140 | 1886251 | recQ & thiC | 112 | CDs037 TPP riboswitch (THI element) (1976113-1976224). |
| 1899755 | 1899879 | CDR20291_1614 & 1615 | 125 | CDR20291_CDs021. Loci present in strain 630 (1989726-1989850). |
| 1935574 | 1935862 | CDR20291_1645 & 1646 | 312 | CDR20291_CDs022. Loci present in strain 630 (2030765-2031076). |
| 1960268 | 1960546 | CDR20291_1668 & 1669 | 279 | CDs039 T-box leader (2055462-2055740). |
| 1971457 | 1971681 | truA2 & argG | 225 | CDs040 T-box leader (2066659-2066883). |
| 2013647 | 2013749 | CDR20291_1718 & cysD | 103 | CDs042 S box leader (2110692-2110794). |
| 2068349 | 2068563 | CDR20291_1771 & 1772 | 215 | CDR20291_CDs023. Loci not present in strain 630. |
| 2123011 | 2123397 | CDR20291_1808 & 1809 | 387 | CD630_n00640 ncRNA IGR (2180693-2181055). |
| 2133184 | 2133305 | CDR20291_1814 & 1815 | 122 | CD630_n00660 (RCd1) (2199358-2199543). |
| 2224501 | 2224856 | CDR20291_1904 & 1905 | 397 | CD630_n00680 (RCd5) (2285913-2286311). |
| 2227154 | 2227277 | CDR20291_1906 & 1907 | 124 | CDR20291_CDs024. Loci not present in strain 630. |
| 2234856 | 2235134 | trpP & CDR20291_1913 | 279 | CDs044 T-box leader (2294385-2294663). |
| 2273973 | 2274204 | argC & CDR20291_1948 | 231 | CDs045 T-box leader (2347170-2347400). |
| 2281478 | 2281581 | CDR20291_1954 & 1955 | 104 | CDR20291_CDs025. Loci not present in strain 630. |
| 2289765 | 2289933 | lysC & CDR20291_1962 | 169 | CDs046 Lysine riboswitch (2368975-2369143) |
| 2308153 | 2308266 | CDR20291_1978 & 1979 | 114 | CDR20291_CDs026. Loci present in strain 630 (2387372-2387485). |
| 2367708 | 2367815 | CDR20291_2022 & bipA | 108 | CDR20291_CDs027. Loci present in strain 630 (2446921-2447028). |
| 2371455 | 2371649 | CDR20291_2024 & thrC | 195 | CD630_n00720 ncRNA IGR (2450669-2450999). |
| 2372114 | 2372358 | CDR20291_2024 & thrC | 245 | CDs048 T-box leader (2451328-2541572). |
| 2514822 | 2515072 | asnC & CDR20291_2145 | 251 | CDs049 T-box leader (2598418-2598668). |
| 2556844 | 2557031 | CDR20291_2173 & 2174 | 188 | CDR20291_CDs028. Loci present in strain 630 (2640435-2640592). |
| 2582664 | 2582815 | CDR20291_2197 & cspD | 152 | Cdi1_9 GEMM RNA motif (2671800-2671951). |
| 2591051 | 2591188 | CDR20291_2206 & 2207 | 138 | CD630_n00820 Antisense CDS (2680182-2680319). |
| 2631113 | 2631213 | grdC & grdB | 101 | CD630_n00830 Antisense CDS (2631113-2631213). |
| 2636494 | 2636691 | grdX & CDR20291_2245 | 198 | CDmisc_RNA_16 glycine (2725664-2725774). |
| 2753450 | 2753648 | glcK & dnaJ | 199 | CDR20291_CDs029. Loci present in strain 630 (2838097-2838295). |
| 2776908 | 2777036 | CDR20291_2372 & 2373 | 129 | CD630_n00840/sCD2862 ncRNA IGR (2861552-2861680). |
| 2801422 | 2801650 | argH & CDR20291_2393 | 229 | CDs051 T-box leader (2886067-2886295). |
| 2825958 | 2826172 | leuS & CDR20291_2411 | 215 | CDs052 T-box leader (2915733-2915947). |

| | | | | |
|---|---|---|---|---|
| 2924278 | 2924462 | CDR20291_2492 & trpS | 185 | CDR20291_CDs030. Loci present in strain 630 (3015643-3015869). |
| 2926387 | 2926642 | mtnN & CDR20291_2495 | 256 | CDs054 T-box leader (3017752-3018007). |
| 2932807 | 2932975 | CDR20291_2500 & 2501 | 169 | CDR20291_CDs031. Loci not present in strain 630. |
| 2934257 | 2934438 | CDR20291_2501 & ileS | 227 | CDR20291_CDs032. Loci not present in strain 630. |
| 2938209 | 2938458 | ileS & CDR20291_2503 | 250 | CDs055 T-box leader (3027704-3027953). |
| 3028772 | 3029044 | asnA & CDR20291_2584 | 273 | CDs056 T-box leader (3114793-3115065). |
| 3051318 | 3051404 | CDR20291_2599 & 2600 | 87 | CD630_n00910 Antisense CDS (3137370-3137456). |
| 3061949 | 3062159 | CDR20291_2606 & 2607 | 211 | CDs057 RNase P (3148001-3148211). |
| 3110462 | 3110730 | CDR20291_2642 & ptsI | 269 | CD630_n00930 ncRNA IGR (3196622-3196979). |
| 3181243 | 3181439 | CDR20291_2688 & 2689 | 197 | CDR20291_CDs033. Loci present in strain 630 (3271493-3271680). |
| 3216981 | 3217190 | CDR20291_2721 & 2722 | 210 | Cdi1_12 GEMM RNA motif (3303255-3303464). |
| 3220424 | 3220558 | CDR20291_2722 & 2723 | 135 | Cdi2_3 c-di-GMP-II (3306644-3306893). |
| 3222189 | 3222409 | CDR20291_2723 & 2724 | 221 | CDR20291_CDs034. Loci present in strain 630 (3308436-3308656). |
| 3356912 | 3357093 | CDR20291_2835 & 2836 | 182 | CDs058 Cobalamin riboswitch (3484931-3485112). |
| 3488192 | 3488331 | CDR20291_2938 & 2939 | 140 | CDs059 Lysine riboswitch (3607770-3607949). |
| 3572842 | 3572939 | CDR20291_2993 & 2994 | 98 | CDR20291_CDs035. Loci not present in strain 630. |
| 3590914 | 3591263 | CDR20291_3010 & smpB | 350 | CDs060 tmRNA (3687389-3687738). |
| 3682298 | 3682472 | dapA1 & asd | 175 | CDs061 Lysine riboswitch (3773053-3773227). |
| 3771564 | 3771679 | CDR20291_3159 & 3160 | 116 | CDs064 RFN element (3681645-3681760). |
| 3830487 | 3830648 | CDR20291_3213 & 3214 | 162 | CD630_33681 (3935442-3935864). |
| 3831015 | 3831348 | CDR20291_3213 & 3214 | 334 | SQ2429 ncRNA IGR (3936132-3936367). |
| 3892780 | 3892959 | cbiP & CDR20291_3258 | 180 | CDs065 cobalamin riboswitch (4027376-4027555). |
| 3896532 | 3896676 | cobT & CDR20291_3262 | 145 | CDR20291_CDs036. Loci present in strain 630 (4031128-4031272). |
| 3986601 | 3986713 | CDR20291_3350 (pilA) & prs | 239 | Cdi2_4 c-di-GMP-II (4105635-4105873). |
| 3986849 | 3987272 | CDR20291_3350 & prs | 424 | CD630_n01120/sCD4107 ncRNA IGR (4106130-4106469). |
| 4014774 | 4015037 | metG & spmB | 264 | CDs066 T-box leader (4136360-4136623). |
| 4084660 | 4084784 | luxS & CDR20291_3437 | 125 | CD630_35980 (4206264-4206459). |
